# Multimodal lesion mapping in affective blindsight reveals dual amygdala and superior temporal sulcus contributions to nonconscious emotion processing

**DOI:** 10.1101/2025.10.13.682195

**Authors:** A.T. Prabhakar, Kavitha Margabandhu, Christilda John Bosco, Rohan Thomas Jepegnanam, Thanusha Prasad, Sowmiya Sampathkumar, Evelyn Sheena Sunderraj, John Davis Prasad, George Abraham Ninan, Deepti Bal, Harshad Vanjare, Anitha Jasper, Pavithra Mannam, Allison M McKendrick, Olivia Carter, Marta I. Garrido

## Abstract

Affective blindsight, the capacity to discriminate emotional stimuli despite bilateral damage to the primary visual cortex (V1) and without conscious awareness, offers a unique model of non-conscious visual processing. Subcortical pathways involving the pulvinar and amygdala have been proposed, but putative cortical contributions remain unclear. We examined 182 patients, including 31 with bilateral V1 lesions. Among these, 15 had cortical visual loss and 7 showed affective blindsight. Using behavioral testing, lesion symptom mapping, and tractography, we found that preserved pulvinar connectivity with both the posterior superior temporal sulcus (STS) and the amygdala is necessary for affective blindsight. These findings provide causal evidence for a multi-route architecture, identifying the pulvinar–STS pathway, alongside the pulvinar–amygdala pathway, as a critical substrate for non-conscious affective processing.

## Introduction

Blindsight refers to preserved visual abilities despite lesions of the primary visual cortex (V1) that abolish conscious sight^1,2^. It spans domains such as motion, orientation, color, and emotion, and manifests as either accurate responses without awareness (Type 1) or vague sensations without full visual perception (Type 2)^3–5^. Affective blindsight describes the capacity to discriminate emotional expressions presented to the blind field, despite lacking subjective awareness^6–11^. Classical models attribute affective blindsight to alternative visual routes that bypass the primary visual cortex, most prominently a subcortical “low road” involving the superior colliculus, pulvinar, and amygdala^6,11–13^. This pathway is supported by neuroimaging and tract-tracing evidence of pulvinar–amygdala connectivity, and by findings that affective stimuli can engage the amygdala without V1 input^10–12,14–16^. In contrast, an alternative view acknowledges the amygdala’s pivotal role in emotion processing but emphasizes a multiple-pathway framework in which cortical–subcortical interactions jointly mediate residual emotional processing ^17^. Given the proposed role of cortical–subcortical interactions in residual emotional processing, the superior temporal sulcus (STS) emerges as a key cortical candidate. The STS is a multisensory cortical hub crucial for integrating visual, auditory, and somatosensory inputs^18–21^. Within it, the posterior bank (pSTS) plays a central role in recognizing dynamic emotional expressions and projects extensively to the lateral amygdala. The pSTS is part of a recently confirmed third visual pathway that links early visual cortex to human middle temporal cortex (motion-sensitive area hMT+) and pSTS and specializes for dynamic emotion, social perception, and biological motion^22–24^. The pSTS, receiving direct pulvinar input^18^, provides a major cortical route for visual information to reach limbic regions, particularly the lateral amygdala for emotional evaluation^25–27^. Although the role of pSTS in affective blindsight remains underexplored, evidence from blindsight patients TN^28^ and MC^29^, showing intact pSTS with activation preceding the amygdala, suggests that affective blindsight may depend on pulvinar-driven pSTS processing. While the pulvinar–amygdala pathway has been anatomically and functionally demonstrated in humans^30^, the role of the pulvinar–pSTS connections in affective blindsight remain unclear. In this study, we examine whether affective blindsight is specifically related to the structural integrity of these pathways, controlling residual geniculostriate input.

## Results

### Behavioral Results

A total of 182 patients with higher-order visual deficits following focal brain lesions were screened for inclusion. Of these, 31 patients were identified to have bilateral lesions of the calcarine cortex (Fig.S1). On clinical assessment, 15 patients subjectively reported complete loss of vision in the absence of any ocular pathology, which was also confirmed by ophthalmological examination. These patients were diagnosed with severe cortical visual impairment and were included for additional behavioral testing for the presence of blindsight with a two-alternative forced choice (2AFC) paradigm. Emotion recognition was assessed using both static photographs and dynamic 3D avatar videos. For static stimuli, participants attempted a five-alternative forced-choice (5AFC) task (happy, sad, fearful, angry, neutral), and for dynamic stimuli, a six-alternative forced-choice (6AFC) task (happy, sad, angry, fearful, surprised, disgusted). Participants who were unable to perform, or who performed poorly on these tasks, were subsequently tested with a 2AFC version. Motion perception was assessed using a random dot kinematogram task requiring discrimination of motion direction (up, down, left, or right). Color perception was evaluated by naming centrally presented discs in 9 universally recognized colors, with a 2AFC format used for those performing below chance. Object recognition was tested with black-and-white line drawings, supplemented by matching tasks when required. Residual face perception was assessed using a jumbled face discrimination task to evaluate the ability to perceive facial configuration. Among the 15 patients tested, 8 exhibited residual visual abilities consistent with blindsight. Seven showed affective blindsight, performing above chance in emotion recognition tasks despite lacking conscious visual perception. One patient exhibited blindsight limited to motion, reaching, and grasping. Clinical characteristics and behavioral performance are summarized in Tables 1 and 2 respectively. All patients had intact pupillary reflexes and no ocular pathology. Despite impaired object, face, and color recognition, each demonstrated affective blindsight alongside varying degrees of preserved motion perception.

**Table 1.**
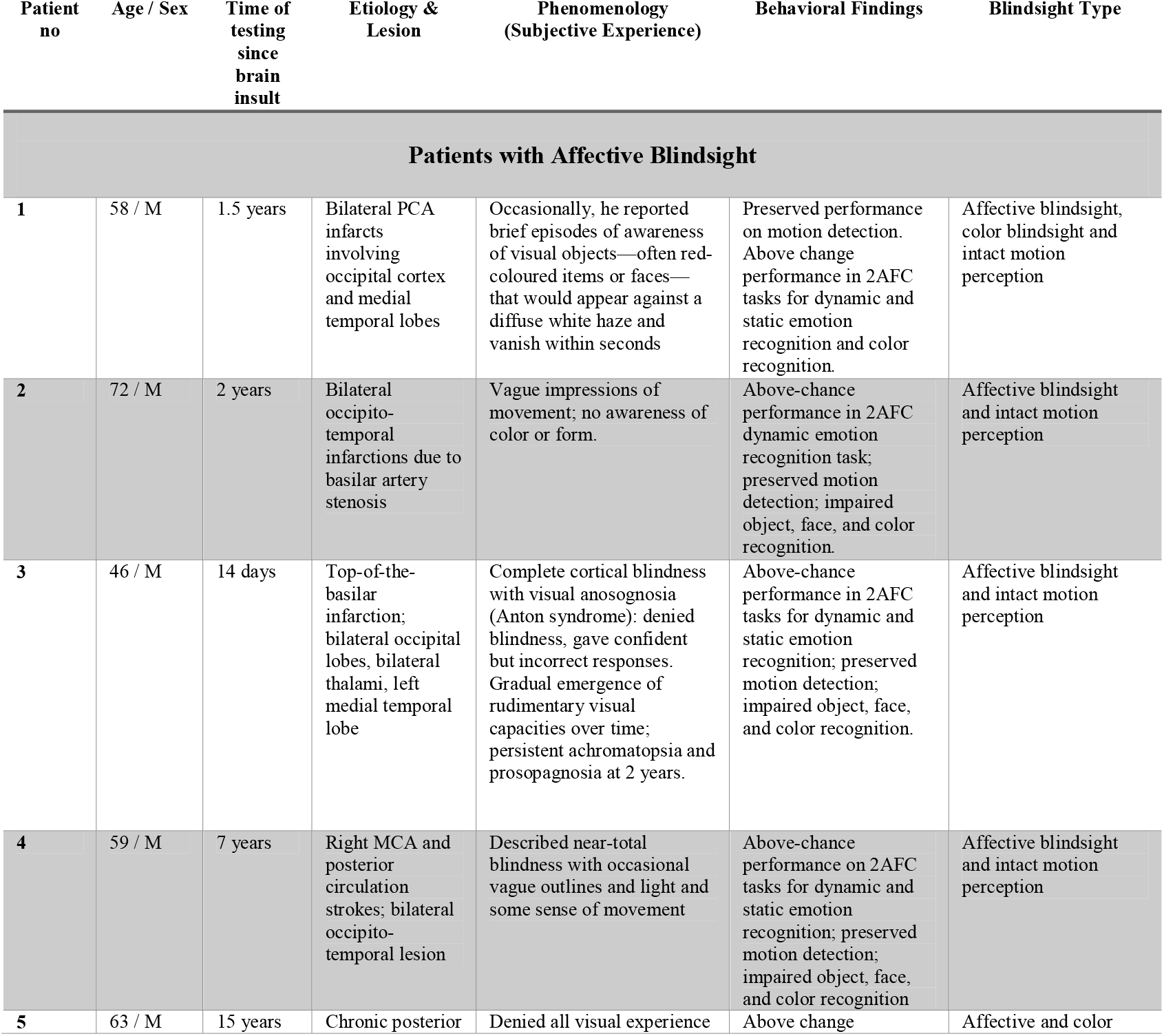
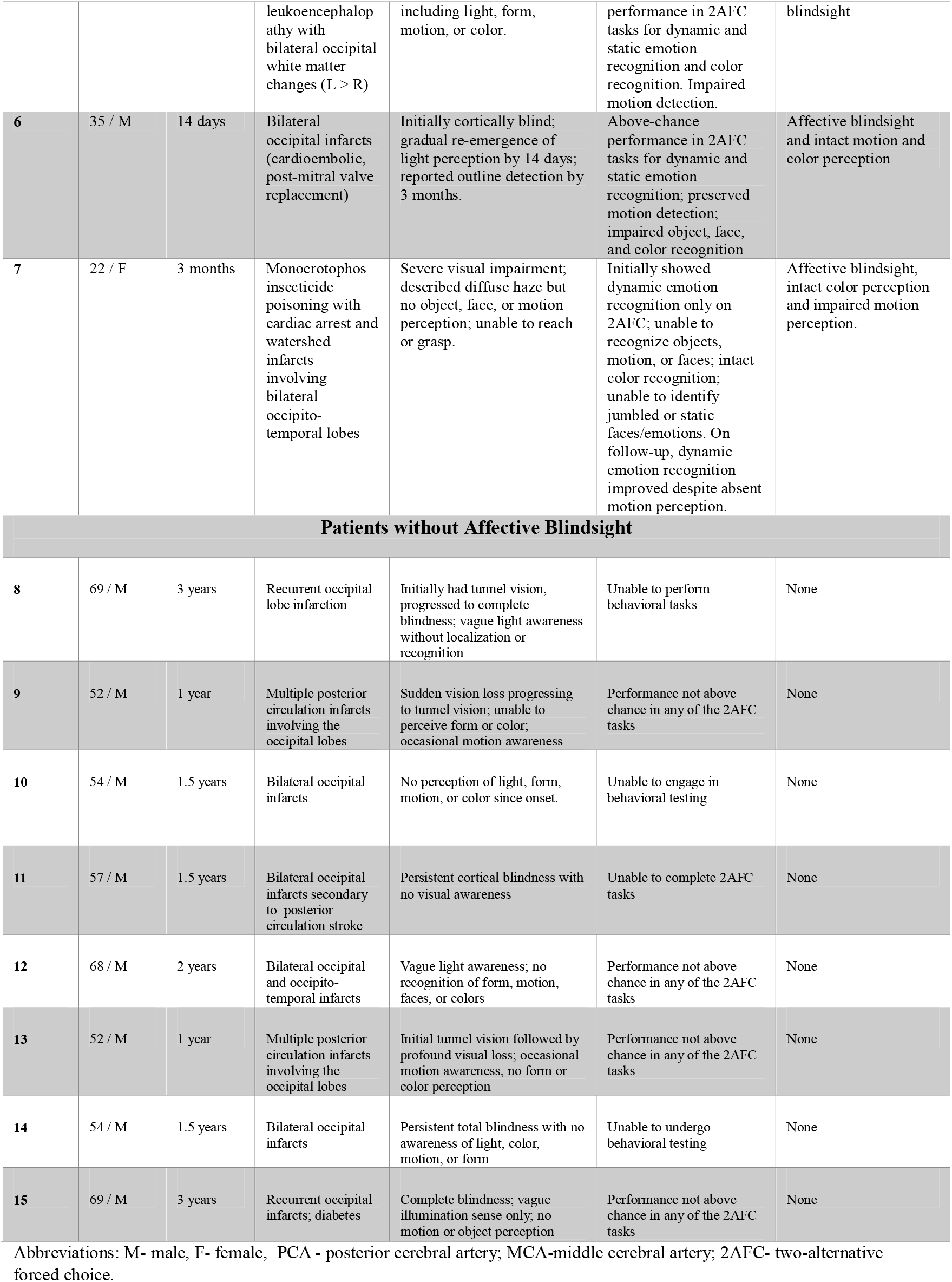
Clinical and Demographic Characteristics of Patients with Affective Blindsight. Abbreviations: M-male, F-female, PCA - posterior cerebral artery; MCA-middle cerebral artery; 2AFC-two-alternative forced choice.

**Table 2.** Performance on Behavioral tasks. Table 2. Performance Accuracy (%) on behavioral tasks in seven patients with cortical visual impairment and affective Blindsight. Accuracy scores are presented for object, color, motion, face, and emotion processing tasks. Recognition tasks refer to direct identification without cues or options, assessing conscious visual perception. 2AFC tasks (two-alternative forced choice) involve selecting one of two options, allowing assessment of non-conscious processing through above-chance performance. Patients were instructed to guess when uncertain. ^*^ - denotes above-chance significance on the binomial test.

| Task | Format | Case 1 | Case 2 | Case 3 | Case 4 | Case 5 | Case 6 | Case 7 |
| --- | --- | --- | --- | --- | --- | --- | --- | --- |
| **Object Naming** | Recognition | 0 | 0 | 0 | 0 | 0 | 0 | 0 |
|  | 2AFC | 16.7 | 20.8 | 6 | 20.8 | 62.0 | 46.7 | 56.0 |
| **Color Naming** | Recognition | 4.2 | 0 | 0 | 0 | 2.1 | 100 | 97.7 |
|  | 2AFC | 70.8* | 54.17 | 47.22 | 66.7 | 100* | — | — |
| **Jumbled Face** | Recognition | 10.0 | 0 | 0 | 10.0 | 0 | 0 | 33.3 |
|  | 2AFC | 55.0 | 45.83 | 41.66 | 56.25 | 52.08 | 48.57 | 62.5 |
| **Static Emotion (Penn)** | Recognition | 0 | 0 | 30 | 0 | 0 | 0 | 0 |
|  | 2AFC | 79.1* | 54.1 | 75.0* | 83.3* | 79.2* | 70.8* | 66.7* |
| **Dynamic Emotion Recognition** | Recognition | 0 | 0 | 11.1 | 16.7 | 22.2 | 27.8 | 0 |
|  | 2AFC | 77.8* | 83.3* | 83.3* | 77.8* | 77.8* | 77.8* | 83.3* |
| **Motion Detection (Dot Task)** | Recognition | 87.5 | 100 | 92.9 | 85.4 | 27.1 | 100 | 0 |
|  | 2AFC | — | — | — | — | 41.7 | — | 41.7 |

### ROI-based analysis confirms Pulvinar’s role in affective blindsight

To explore the neuroanatomical correlates of affective blindsight, we first conducted a univariate analysis comparing lesion involvement in predefined regions of interest (ROIs) between patients with (N = 7) and without (N = 8) affective blindsight. The ROIs included the pulvinar, lateral geniculate nucleus (LGN), human MT+ (hMT+), posterior superior sulcus (pSTS), amygdala, and calcarine cortex. ROIs were delineated using established atlases in MNI space and binarized for lesion overlap. Lesion status (present/absent) for each ROI was derived from patient-specific lesion maps, which were manually traced directly in MNI space from clinical MRI scans, using DWI for acute strokes and T2-FLAIR for chronic lesions and independently verified by two neurologists. Group comparisons for each ROI were performed using Fisher’s exact test, and calcarine lesion volume (voxel count) was compared using an independent samples t-test (see Table 3).

**Table 3.**
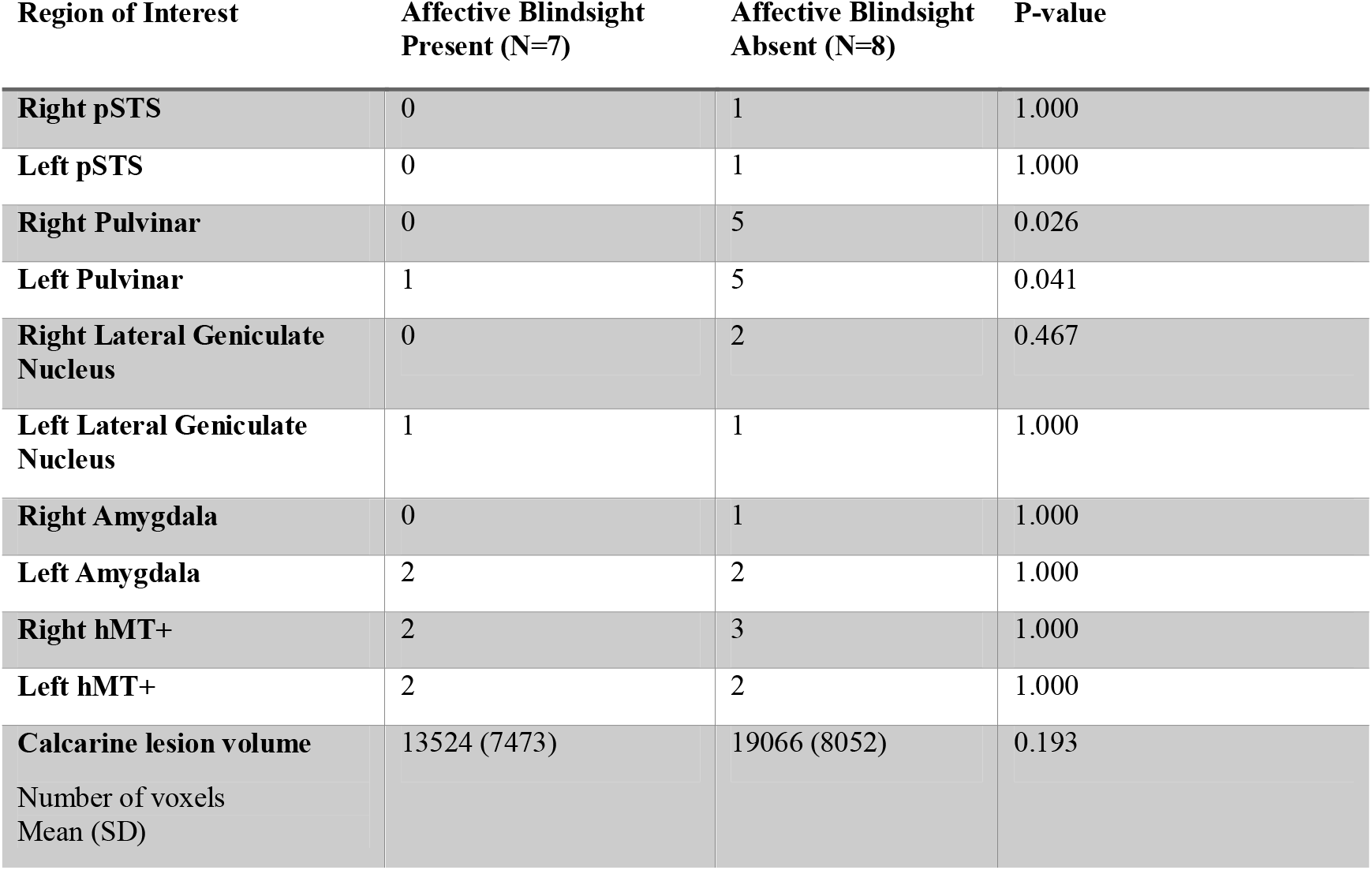
Patients with lesions involving ROIs stratified by presence or absence of Affective Blindsight. Table 3. Univariate Analysis of Lesion Involvement and Presence of Affective Blindsight This table presents a univariate comparison of lesion involvement across predefined regions of interest (ROIs) in patients with and without affective blindsight. For each ROI, the number of patients with lesions in that region is shown, stratified by the presence or absence of affective blindsight (N = 7 and N = 8, respectively). Categorical comparisons were conducted using Fisher’s exact test. Mean calcarine lesion volume (in voxels) was compared using an independent samples *t*-test. Significant associations were observed for the right and left pulvinar, suggesting their potential role in supporting non-conscious affective processing. **Abbreviations:** pSTS – posterior superior temporal sulcus; hMT+ – human middle temporal complex (motion-sensitive area MT/V5); SD – standard deviation.

Significant group differences were observed for bilateral pulvinar. Lesions to the right pulvinar were more frequently observed in patients without affective blindsight (5/8) than those with blindsight (0/7), yielding a statistically significant association (*p* = 0.026). Similarly, the left pulvinar was lesioned in 5 of 8 blindsight-absent patients, compared to only 1 of 7 blindsight-present patients (*p* = 0.041). No significant group differences were found for other ROIs, including the amygdala, lateral geniculate nucleus (LGN), posterior superior temporal sulcus (pSTS), or human middle temporal area (HMT). Although patients without blindsight exhibited numerically greater calcarine lesion volumes (mean = 19,066 voxels, SD = 8,052) compared to those with blindsight (mean = 13,524 voxels, SD = 7,473), this difference did not reach statistical significance (*p* = 0.190). Together, these findings indicate that sparing of the right pulvinar is critical for preserving non-conscious affective processing in patients with bilateral occipital damage, providing confirmatory evidence for the pulvinar’s critical role in non-conscious emotion processing. (see **Figure 1.A**).

**Figure. 1.**
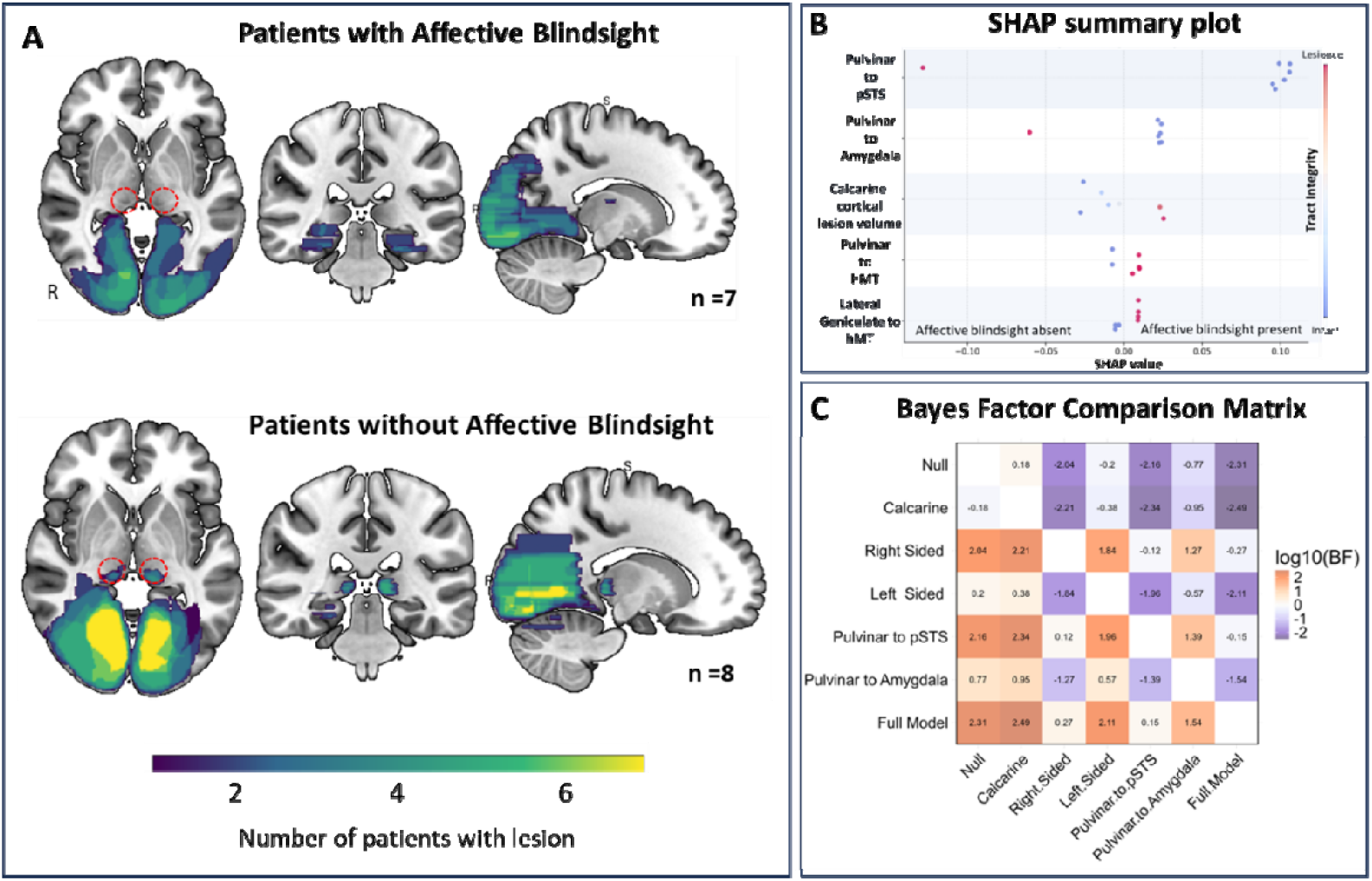
Pulvinar and its tracts to amygdala and pSTS predict affective blindsight. **(A)** Shows lesion overlap maps of patients with cortical visual impairment with affective blindsight (1^st^ row) and those without (2^nd^ row). Lesion maps are displayed on the MNI152 template brain. Color intensity reflects the number of patients with damage at each voxel, with warmer colors (yellow) indicating higher overlap. Pulvinar involvement (location highlighted by red rings in the axial slices) was observed more commonly in patients without affective blindsight. **(B)** Shows the summary of SHAP (SHapley Additive exPlanations) analysis on the logistic regression model illustrating the contribution of each tract of interest (TOI) to the classification of affective blindsight. Each dot represents a patient’s SHAP value for a given TOI; dots to the right indicate features contributing more to the presence of affective blindsight, while those to the left contribute to its absence. Dot color reflects tract integrity—blue for intact, red for lesioned. The plot shows that the integrity of the Pulvinar–STS and Pulvinar–Amygdala pathways contributed to the prediction of affective blindsight, while lesions in these tracts was associated with its absence. **(C)** Shows Pairwise Bayes Factor Comparison Matrix (log□□ scale) comparing the explanatory power of seven competing logistic regression models incorporating lesion status in key subcortical tracts—namely, the pulvinar–amygdala and pulvinar–posterior superior temporal sulcus (pSTS) pathways. Each cell in the heatmap shows the log□□-transformed Bayes Factor (BF), comparing the model in the row (numerator) against the model in the column (denominator). Positive values (orange) indicate greater support for the row model; negative values (purple) favour the column model; values near zero (white) indicate negligible evidence in either direction. Model comparisons are based on marginal likelihoods estimated using bridge sampling. The symmetric color gradient ranges from purple (log□□BF = –2.5) to orange (log□□BF = +2.5), centered at zero (white). **Abbreviations:** SHAP: SHapley Additive exPlanations; pSTS-posterior superior temporal sulcus; hMT - human middle temporal area (motion sensitive area)

### Integrity of right tracts from pulvinar to amygdala and pulvinar to pSTS predict affective blindsight

To investigate the contribution of white matter pathways projecting from the pulvinar to the amygdala and extrastriate temporal cortex in affective blindsight, we conducted a tract-of-interest (TOI) analysis focusing on the pulvinar–amygdala and pulvinar–pSTS pathways. We performed disconnectome mapping using the BCBtoolkit, which estimates likely tract disruptions by seeding patient lesion maps into normative tractography data from healthy controls. This approach allowed us to infer structural disconnection beyond the lesion site and evaluate TOI involvement. Tracts intersecting lesioned or disconnected voxels were coded as disconnected, and all TOIs were binarized as disconnected or spared. A logistic regression with L2 regularization was performed to predict affective blindsight. Predictors included binary indicators of tract disconnection for right and left pulvinar–amygdala, pulvinar–pSTS, and LGN–hMT+ pathways, along with calcarine lesion volume (voxel count) as a covariate. (see **Table 4**). The results revealed that damage to the right pulvinar–amygdala and right pulvinar–pSTS tracts was strongly associated with reduced likelihood of affective blindsight (*p* = 0.01 for both), with odds ratios of 0.43 (95% CI: 0.34–0.56). In contrast, left-sided homologous tracts and lateral geniculate pathways were not significant predictors (see Table 4). Lesion volume in the calcarine cortex was also not associated with affective blindsight in this model (*p* = 0.5475). These findings highlight the critical role of two distinct right-sided subcortical pathways, specifically from the pulvinar to the amygdala and pulvinar to pSTS, in supporting non-conscious affective processing in the absence of visual awareness.

**Table 4.** Multivariate Ridge Regression Model Predicting Affective Blindsight from Tract Integrity. **Table 4** Ridge-Regularized Logistic Regression Identifying Lesioned Tract Predictors of Affective Blindsight. The table summarizes the results of a multivariate logistic regression with L2 (Ridge) regularization assessing the association between tract-specific lesions and the absence of affective blindsight. Lesions and disconnectomes were mapped onto tractography-defined regions of interest (TOIs), including pulvinar–amygdala and pulvinar–posterior superior temporal sulcus (pSTS) pathways. Calcarine lesion volume was included as a covariate. Significant associations were observed for right pulvinar– amygdala and right pulvinar–pSTS tracts, indicating their key role in affective blindsight. **Abbreviations:** pSTS = Posterior Superior Temporal Sulcus, LGN = Lateral Geniculate Nucleus, hMT+ – human middle temporal complex (motion-sensitive area MT/V5), CI = Confidence Interval. ^*^-denotes statistically significant values.

| Feature | Mean Coefficient | Std. Dev. | Z-score | p-value | Log Odds | Odds Ratio (95% CI) |
| --- | --- | --- | --- | --- | --- | --- |
| **Right Pulvinar to Amygdala** | -0.854 | 0.148 | -2.565 | 0.01* | -0.854 | 0.426 (0.339–0.589) |
| **Left Pulvinar to Amygdala** | 0.214 | 0.336 | 0.411 | 0.707 | 0.214 | 1.238 (0.728–2.379) |
| **Right Pulvinar to pSTS** | -0.854 | 0.148 | -2.565 | 0.01* | -0.854 | 0.426 (0.339–0.589) |
| **Left Pulvinar to pSTS** | -0.533 | 0.261 | -1.070 | 0.306 | -0.533 | 0.587 (0.358–0.952) |
| **Right LGN to hMT+** | 0.112 | 0.235 | 0.213 | 0.837 | 0.112 | 1.118 (0.713–1.699) |
| **Left LGN to hMT+** | -0.127 | 0.264 | -0.235 | 0.857 | -0.127 | 0.881 (0.520–1.413) |
| **Calcarine** | -0.339 | 0.242 | -0.644 | 0.540 | -0.339 | 0.713 (0.429–1.068) |

To compare the contribution of pulvinar–amygdala versus pulvinar–pSTS pathways to affective blindsight, we conducted a SHAP (SHapley Additive exPlanations) analysis on the fitted logistic regression model. SHAP provides a unified, game-theoretic approach to explain individual predictions by estimating the marginal impact of each predictor on the model’s output^31^. To interpret the contribution of each tract of interest (TOI) to the prediction of affective blindsight, we generated SHAP summary plot (**Figure 1.B**). Each dot on the plot represents a patient’s SHAP value for a given TOI, indicating how much that feature contributed to the model’s prediction for that individual. Dots farther to the right contributed more to the model predicting the presence of affective blindsight, while those to the left contributed to its absence. Dot color indicates tract integrity—blue for intact, red for lesioned. Our analysis revealed that both the pulvinar–amygdala and pulvinar–pSTS pathways significantly influenced the model’s predictions. Specifically, preservation of these tracts contributed more to the presence of affective blindsight, while lesions in these pathways contributed to the absence of affective blindsight.

### Bayesian Model Comparison Reveals Preservation of Right-Lateralized Subcortical Pathways Supports Affective Blindsight

To complement and validate our findings, we used Bayesian logistic regression, which handles small samples and correlated predictors while enabling inference and model comparison via Bayes Factors. Models were estimated using Markov Chain Monte Carlo sampling with weakly informative priors. Marginal likelihoods were estimated using bridge sampling, a method that combines samples from the posterior and a simpler proposal distribution to yield stable and accurate estimates of model evidence. To assess the contribution of each tract, we computed log□□-transformed Bayes Factors (BFs) by comparing models with and without each predictor. This allowed us to quantify the relative evidence for the inclusion of each tract in predicting affective blindsight performance (**Figure 1.C**). We compared seven competing models of tract involvement. These included: (1) a Null model with no lesion-defined tract predictors, (2) a Calcarine-only model (*Calcarine*) including only calcarine damage, (3) a Pulvinar–amygdala pathway model (*Pulvinar to Amygdala*) incorporating bilateral pulvinar–amygdala interactions, (4) a Pulvinar–pSTS pathway model (*Puvinar to pSTS*) comprising bilateral pulvinar–pSTS interactions, (5) a Left-Sided Pathways model (*Left sided*) with left pulvinar–amygdala and pulvinar–pSTS tract interactions, (6) a Right-Sided Pathways model(*Right sided*) with corresponding right hemisphere interactions, and (7) a full model (*Full Model*) incorporating calcarine damage along with all main effects and pairwise interactions among the four lesion-defined tracts. Our results showed that both the pulvinar–pSTS and pulvinar–amygdala pathways contribute to affective blindsight, with stronger effects associated with the pulvinar–pSTS tract and right hemisphere involvement (**Figure 1C**). Although the Full Model, which included all main effects and interactions, achieved the highest overall support, the pulvinar–pSTS model performed nearly as well and offers a more parsimonious account of the data (**Figure 1C**). By contrast, the Calcarine-only model performed worse than the Null model, providing no evidence for a role of primary visual cortex damage in residual affective processing and not supportive of the view that blindsight can be explained by ‘islands’ of spared vision within V1.

### Disconnectome mapping shows Pulvinar to amygdala and Pulvinar to Posterior superior temporal sulcus tract integrity supports affective blindsight

We conducted a multimodal lesion analysis to identify the structural and functional neural correlates of affective blindsight. Using manually delineated lesion maps for each patient, we derived both indirect functional and structural connectivity measures. For lesion-network mapping, each lesion map served as a seed to generate functional connectivity maps based on a normative resting-state fMRI connectome, identifying regions functionally connected to the lesion site in healthy individuals. To assess structural disconnection, we used a normative diffusion-weighted imaging (DTI) dataset to generate patient-specific disconnectome maps, estimating the probability of white matter tract disruption caused by each lesion. These three lesion-derived modalities—lesion map, lesion network map, and disconnectome map—were then analyzed using multivariate lesion–symptom mapping. We applied both support vector regression-based lesion–symptom mapping (SVR-LSM) and Bayesian Lesion–Deficit Inference (BLDI) to identify associations between these neural features and the presence of affective blindsight. Of these, only the disconnectome mapping using BLDI yielded significant results (**Figure 2.A**). Crucially, preservation of right-lateralized pulvinar - amygdala tracts and pulvinar– posterior superior temporal sulcus projections were predictive affective blindsight. In the Pulvinar–pSTS projections, particularly the Arnold’s tract (AR type, right hemisphere) was found to be associated with affective blindsight. Moreover, the preservation of long-range associative tracts, including the middle longitudinal fasciculus, and arcuate fasciculus, appeared to support higher-order integration between occipitotemporal visual areas and frontal-limbic networks implicated in affective salience detection.

**Figure 2:**
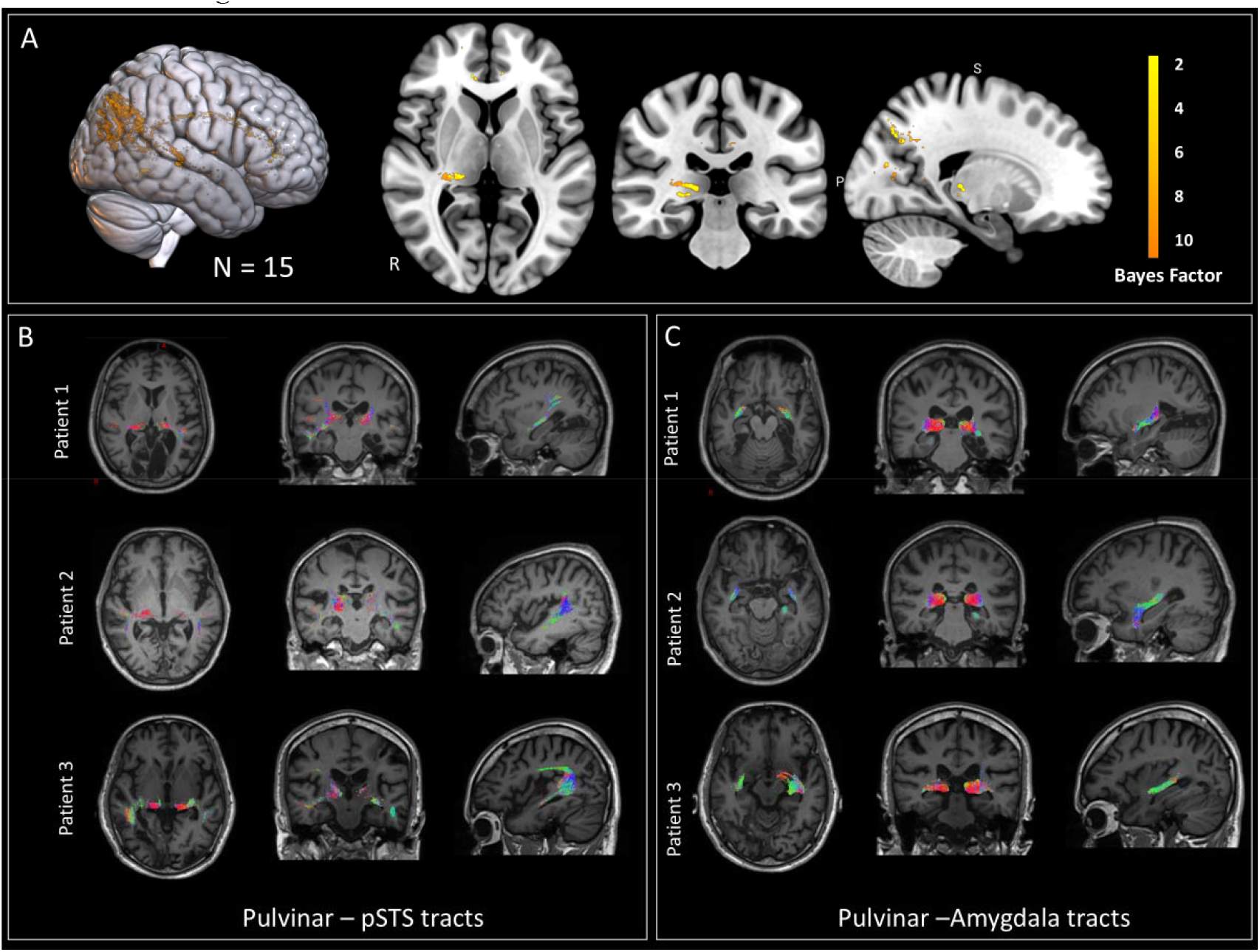
Bayesian lesion deficit inference and diffusion tractography in patients with affective blindsight. (A) Rendered MNI152 template brain and orthogonal brain slices show results from the Bayesian Lesion– Deficit Inference (BLDI) with lesion derived disconnectomes as input to identify the white matter tracts associated with the presence of affective blindsight. Clusters (Bayes factor > 2) indicate preserved white matter pathways predictive of affective blindsight, including the pulvinar–STS and pulvinar–amygdala tracts, as well as long-range associative pathways such as the middle longitudinal fasciculus and arcuate fasciculus (B) Results from local diffusion tractography analysis in three patients with affective blindsight showing preserved pulvinar– posterior superior temporal sulcus (pSTS) tracts. Each row corresponds to a different patient; columns 1–3 show axial, coronal, and sagittal views, respectively, with the reconstructed pulvinar–STS tract overlaid on the patient’s T1-weighted MRI in MNI space. (C) Corresponding results for the pulvinar–amygdala tracts. Tracts are color-coded by principal diffusion direction: red = left–right, green = anterior–posterior, and blue = superior–inferior.

### Diffusion tractography results confirms preservation of pulvinar-amygdala-STS white matter tracts in patients with blindsight

To confirm the integrity of the identified tracts in patients with affective blindsight, we conducted tractography in three patients who returned for follow-up (Patients 1–3, all with preserved motion perception). Both global and local probabilistic tractography approaches revealed reduced overall fiber count and apparent fiber density (AFD) relative to matched controls (Tables S1–S4). Analysis of the tracts of interest showed no group differences in pulvinar–amygdala connectivity (Left: β = 0.02, 95% CI = −0.32–0.36, p = 0.904; Right: β = 0.10, 95% CI = −0.23–0.43, p = 0.531), whereas pulvinar–pSTS connectivity was significantly reduced in patients (Left: β = −3.61, 95% CI = −5.59–−1.63, p = 0.0004; Right: β = −2.64, 95% CI = −4.47–−0.82, p = 0.0048). Importantly, both pulvinar–pSTS and pulvinar–amygdala tracts were still identifiable in all three patients with affective blindsight (Figs. 6, 7).

## Discussion

Our findings provide convergent evidence from multiple analytic approaches, showing that affective blindsight relies on both subcortical and cortical visual–limbic pathways, consistent with a “multiple-roads” model of non-conscious emotion processing. Beyond the amygdala, the pulvinar–pSTS pathway emerges as a key cortical route linking dynamic emotional cues to limbic processing. In the region-of-interest analysis aimed at identifying anatomical regions associated with affective blindsight, we found that none of the patients with pulvinar lesions, particularly in the right hemisphere, showed sparing of emotion processing. These findings provide additional confirmatory evidence for the critical role of the pulvinar in affective blindsight^12,32–34,34–36^. Further, our tract-of-interest analysis revealed that lesions to right-sided pulvinar projections, specifically the pulvinar–amygdala and pulvinar–posterior superior temporal sulcus (pSTS) pathways were significantly associated with the absence of residual emotion processing. Bayesian model comparisons further reinforced these findings, showing that models incorporating right side and pulvinar–pSTS and pulvinar–amygdala pathways provided the strongest explanatory power, while cortical-only models were insufficient. These results indicate that, in addition to the amygdala, the pSTS plays a significant role in affective blindsight^35,37,38^. This pattern of findings aligns more closely with the multiple-roads model, wherein parallel and interacting subcortical and cortical pathways—rather than a single dedicated “low road”—jointly support non-conscious emotion processing^14,17,33,39,40^. Converging evidence from Bayesian lesion-deficit inference based on the disconnectome maps also showed involvement of the pulvinar and its projections to both the amygdala and pSTS^41^. Additional associations between affective blindsight and the middle longitudinal fasciculus (MDLF) as well as posterior segments of the arcuate fasciculus suggest that, beyond early subcortical relays, a broader temporo-parietal-limbic network contributes to non-conscious emotion processing. The pSTS, hMT+, and MDLF, all implicated in blindsight, form part of the proposed third visual pathway, specialized for motion perception, dynamic emotional expressions, and social perception^22,24,42^. We propose that this pathway, optimized for rapid action, plays a central role in affective blindsight and non-conscious affective processing^22,24,42^. In our prior work with focal brain lesion patients, we demonstrated a double dissociation between dynamic and static emotion recognition, providing causal evidence for this third pathway^24,43^. Specifically, lesions involving either the pSTS or the pulvinar (even in the absence of direct pSTS involvement) impaired dynamic emotion recognition, underscoring the critical role of both structures and their connecting white matter tracts^24^. Patients with affective blindsight consistently performed better on tasks involving dynamic compared to static emotional faces, reinforcing the view that the residual visual system is preferentially tuned to motion-based affective cues—a capacity likely mediated by the pSTS, which encodes dynamic emotional and biological motion signals^19,21,22,44^. The pulvinar, receiving direct input from motion-sensitive neurons in the superior colliculus, may act as a conduit for rapidly transmitting this information to the STS, bypassing damaged striate cortex^12,14,39,45,46^. Taken together, these findings support the conclusion that the pulvinar–pSTS pathway provides a crucial subcortical input to the third visual pathway and underlies the dynamic emotion processing observed in affective blindsight, thereby contributing fundamentally to non-conscious affective perception. Tractographic data from a subset of patients with affective blindsight(n = 3) who were available for follow up provided further support for this model, with all three individuals demonstrating partially preserved pulvinar–pSTS and pulvinar–amygdala tracts relative to age-matched controls. Our findings support a V1-independent visual system for affective and social cues, relying on a subcortical–temporal network involving the pulvinar, pSTS, and amygdala. This provides a structural basis for preserved affective perception without visual awareness and highlights the multidimensional nature of blindsight. From a clinical perspective, it is important to recognize that bilateral calcarine lesions are most commonly caused by infarction within the posterior cerebral artery (PCA) territory, which may variably involve the pulvinar. In such cases, key regions including hMT+, pSTS, and the amygdala are often spared, as they fall within the territories of the middle cerebral artery (MCA) and anterior choroidal artery, respectively. This vascular distribution implies that pulvinar sparing may serve as a critical determinant for the presence of affective blindsight. Clinically, this has important prognostic implications, such that patients with cortical visual impairment and an intact pulvinar may demonstrate more favorable long-term outcomes, namely residual vision, albeit non-conscious.

## Limitations

Several methodological limitations warrant consideration. First, due to the hospitalized patient population, behavioral testing was necessarily brief and constrained in scope, as individuals with severe cortical visual impairment often exhibit reduced attentional capacity and limited tolerance for prolonged assessments. Second, the ROI-based analyses relied on standard atlas-defined regions rather than patient-specific functional localizers; however, given the extent of cortical damage, these differences are likely of limited consequence. Third, tractography analysis was conducted in only three patients, limiting statistical power and generalizability. Additionally, the sample size for lesion mapping and BLDI was relatively small, which may have constrained the detection of more subtle lesion–behavior relationships. However, given the rarity of blindsight, this represents a substantial and informative cohort. Lastly, both lesion mapping and BLDI are subject to inherent challenges, including image normalization errors, segmentation variability, and anatomical differences across individuals, all of which may impact spatial accuracy and interpretability. However, the use of a multimodal approach helps to mitigate some of these limitations.

## Conclusion

This study provides evidence that affective blindsight is supported by right-lateralized pathways linking the pulvinar with both the amygdala and pSTS. These findings support a multiple-roads model of non-conscious emotion processing that extends beyond the classical amygdala route and highlight the pSTS’s important role in enabling emotional perception without visual awareness.

## Methods

### Participants

Patients were prospectively recruited from the Christian Medical College (CMC) Vellore Brain Lesion Registry between January 2022 and April 2025. Inclusion criteria comprised: (1) radiologically confirmed focal cortical lesions involving temporal, parietal or occipital lobes; (2) availability of high-quality multiplanar MRI sequences including DWI and FLAIR; (3) clinical stability permitting behavioral testing. Exclusion criteria eliminated: (1) impaired consciousness or attention; (2) primary ocular pathology; (3) communication deficits; (4) diffuse or non-cortical lesions. From an initial pool of 182 patients with higher-order visual dysfunction, 31 demonstrated bilateral V1 lesions on MRI, of whom 15 met criteria for cortical visual loss. Seven patients exhibited affective blindsight on formal testing. As the participants were visually impaired, verbal informed consent was obtained directly from them, and written informed consent was provided by a close relative, in accordance with the protocols approved by the CMC Vellore Institutional Review Board (IRB Min No 11542).

### Behavioral Paradigms

Behavioral tasks were implemented in PsychoPy v3.0 and presented binocularly on a 1920×1080 monitor (60 Hz refresh rate) at a 70 cm viewing distance. As all patients included in the study had cortical visual impairment, the monitor location and stimulus display area were indicated by guiding each participant’s hand to the screen center, after which they were instructed to maintain fixation at that position. Eye movements and attention lapses were monitored throughout. Patients performing poorly on the primary tasks underwent two-alternative forced choice (2AFC) testing to confirm blindsight, while those with residual weakness who could not be tested in the laboratory were assessed at the bedside using a standardized laptop-based paradigm.

### Static facial expressions

To assess emotion recognition, we first used static facial images from the Penn Emotion Recognition Task (ER-40), a standardized set of 40 color photographs depicting individuals of diverse ethnic backgrounds expressing four basic emotions—happiness, sadness, anger, and fear—along with a neutral expression^47^. Each image was presented centrally, occupying a visual angle of 10 × 12 degrees. Participants identified the emotion via a five-alternative forced choice (5AFC: happy, sad, fearful, angry, or neutral) as the response options were read aloud in sequence until a response was given. If no response was made within 30 seconds despite attention to the image, the trial was terminated and the next began after a 1000 ms interstimulus interval. Among neurotypical controls (N = 68), the mean accuracy was 85% (SD = 7%). A cutoff of 71% (2 SDs below the mean) was used to indicate impaired performance. Fifteen participants who failed to complete the 5AFC task in view cortical visual loss were subsequently assessed using a simplified two-alternative forced choice (2AFC) version with the same stimuli.

### Dynamic facial expressions

To assess dynamic emotion recognition, facial expression stimuli were created using the FACSHuman plugin for the MakeHuman software^48^. FACSHuman allows manipulation of nearly all Facial Action Units (AUs) defined in Ekman’s Facial Action Coding System (FACS) through a 3D modeling interface. Using this tool, we generated three avatars (male, female, and gender-neutral), and created six video clips displaying the emotions: happiness, sadness, anger, fear, surprise, and disgust. Each video began with a neutral face and gradually intensified the target emotion to 80% over 1.5 seconds^24^. Videos were looped and displayed centrally on the screen at a visual angle of 10° × 12°. Participants identified the emotion using a six-alternative forced choice (6AFC: happy, sad, angry, fearful, surprised, or disgusted) via verbal response. As the participants were visually impaired the options were read aloud sequentially until a response was selected. Each trial was displayed till the participant responded and was followed by an interstimulus interval of 500 ms. If the participant gave no response within 30 seconds despite paying attention, the trial was terminated. Each participant completed one block of 18 trials. Normative data from a neurotypical sample (N = 73) showed a mean accuracy of 66% (SD = 11%). A cutoff score of 44% (2 SDs below the mean) was used to indicate impaired performance. For the fifteen participants with bilateral calcarine lesions who were unable to perform the 6AFC task or scored below the cutoff completed a simplified two-alternative forced choice (2AFC) version of the task, comprising 48 trials.

### Motion direction discrimination task

Motion perception was assessed using a global motion direction discrimination task based on a random dot kinematogram (RDK). Stimuli were generated using the Dots component in PsychoPy, following the design of direction discrimination paradigms used in stroke populations by Vaina et al^49^. The display consisted of 158 white dots (luminance: 79.2 cd/m^2^, subtending 4 arcmin each) presented on a grey background within a circular aperture of 10° diameter, centered on the screen. Dot density was 2 dots/deg^2^, with a constant speed of 3°/s.

Each dot lasted 22 frames, after which a new dot was generated at a random position. All dots moved coherently (100% coherence) in one of four cardinal directions (90°, 180°, 270°, or 360°), randomly selected per trial. The stimulus remained visible until the participant responded using a four-alternative forced choice (4AFC: up, down, left, or right). Each participant completed 24 trials. Participants with bilateral calcarine lesions who were unable to complete the 4AFC task or scored below chance underwent a simplified two-alternative forced choice (2AFC) version of the motion perception task. This version included 48 trials and used the same RDK parameters but with only two motion directions per trial.

### Color naming Task

This task assessed basic color perception by requiring participants to identify and name the color of a circular disc presented at the center of the screen^50^. Stimuli consisted of nine universally recognized colors across the Indian subcontinent: black, blue, brown, green, pink, purple, red, yellow, and white. Colored discs were created using PsychoPy and subtended a visual angle of 10°. Each participant completed 36 trials, responding verbally from a fixed set of nine color options. Fifteen participants who were unable to perform the 9-alternative forced choice (9AFC) task or scored below chance underwent a simplified two-alternative forced choice (2AFC) version consisting of 48 trials, using the same stimulus set but limited to two color choices per trial.

#### Object naming task

This task involved naming 50 simple black-and-white line drawings of everyday objects, each presented centrally on the screen at a size of 10° × 12° visual angle. The task was adapted from Newcombe et al.’s work on patients with brain lesions^51^. Participants gave verbal responses, which were scored as correct or incorrect by the examiner. If no response was made within 30s despite attention to the stimulus, the trial was terminated and followed by a 500ms interstimulus interval. Fifteen participants who showed impaired performance on the object naming task underwent a two-alternative forced choice (2AFC) version of the task consisting of 50 trials. In addition, object and shape matching tasks were administered to evaluate for visual agnosia.

### Jumbled face recognition task

To assess coarse face perception in cortically blind patients who were unable to perform object recognition, a two-alternative forced choice (2AFC) jumbled face task was administered^52^. The task comprised 20 trials: 10 normal faces and 10 jumbled faces. In the jumbled condition, facial features (eyes, nose, mouth) were rearranged within the facial outline while preserving overall shape and contrast. On each trial, a single face stimulus was presented centrally, and participants were asked to verbally indicate whether the face was normal or jumbled. This allowed evaluation of residual facial form discrimination despite impaired object-level vision.

### Behavioural Data Analysis

Accuracy (percentage of correct responses) was calculated separately for each experimental condition. For participants able to perform the regular behavioral tasks, scores were z-transformed using normative values from neurotypical controls^24^, whereas performance on 2AFC tasks was assessed with a binomial test to determine responses above chance.

### Neuroimaging Protocols

#### MRI Data Acquisition

All patients underwent clinical MRI on a 3T Siemens Skyra scanner. The imaging protocol included T2-FLAIR, diffusion-weighted imaging (DWI; b = 1000 s/mm^2^), and susceptibility-weighted imaging (SWI). Lesions were manually delineated from these sequences and directly mapped onto the MNI template using MRIcroGL^53^. Three patients with affective blindsight, whose vision had partially recovered and were available for follow-up, underwent additional detailed neuroimaging. This included high-resolution anatomical scans acquired using a T1-weighted MPRAGE sequence (TR = 2300 ms, TE = 2.98 ms, flip angle = 9°, voxel size = 1 mm^3^ isotropic, FOV = 256 mm, 176 slices), which were used for lesion delineation, spatial normalization, and co-registration with diffusion data. Diffusion tensor imaging (DTI) was performed using a single-shot spin-echo EPI sequence with 64 diffusion-weighted directions (b = 1000 s/mm^2^) and an isotropic resolution of 2.4 mm^3^.

#### Lesion Mapping

Clinical MRI data, including T2-weighted, T2-FLAIR, T1-weighted, and DWI sequences, were reviewed to delineate lesions. For acute ischemic strokes, DWI was primarily used, while T2-FLAIR was preferred for chronic lesions. Lesions were manually traced in Montreal Neurological Institute (MNI) space using MRIcroGL^53^ and saved a binary nifty files. Mapping was performed in 3D using axial, coronal, and sagittal views by an experienced neurologist (ATP), and independently cross-verified by a second neurologist. Lesion boundaries were finalized by consensus.

### Region and tract of interests (ROI/TOI) analysis

To examine the relationship between lesion location and the presence of affective blindsight, a region-of-interest (ROI)–based analysis was conducted. The selected ROIs included the bilateral pulvinar, lateral geniculate nucleus (LGN), posterior superior temporal sulcus (pSTS), human motion-sensitive area (hMT+), amygdala, and bilateral calcarine cortex (Fig. 1). All ROIs were anatomically defined using segmentations from the FreeSurfer atlas, except for hMT+, which was defined using Wang’s probabilistic visual atlas, thresholded at 80% probability^54–57^. Tracts of interest (TOIs) were defined for the following pathways in both hemispheres: pulvinar–MT, LGN–MT, pulvinar–pSTS, and pulvinar–amygdala. Masks for the pulvinar–MT, LGN–MT, and pulvinar–pSTS tracts were obtained from published literature^41,58^. The pulvinar–amygdala tracts, which have also been described in prior studies, were generated in-house using normative diffusion data^30^. All tract masks were binarized and overlapped with lesion maps and thresholded disconnectome maps to assess tract involvement. Involvement of each ROI and TOI (excluding calcarine cortex) was coded as a binary variable (lesioned or spared). The volume of lesion overlap with the bilateral calcarine cortex—given its established role in visual awareness—was included as a continuous covariate. Initial comparisons were conducted to assess associations between individual region-of-interest (ROI) involvement and the presence of affective blindsight, defined as above-chance performance in the dynamic emotion recognition task and treated as a binary outcome (present/absent). Depending on cell sizes, either Fisher’s exact test (for expected counts <5) or chi-square test was used. To account for the influence of multiple lesion sites and regress out the effect of the calcarine cortex, we conducted a multivariate analysis to evaluate the independent contribution and interaction of each ROI and TOI. ROI and TOI involvement was coded as binary variables, with 1 indicating a lesioned region/tract and 0 indicating a spared one. First, we employed logistic regression with regularization to address high multicollinearity among predictors and the limited sample size. Both L1 (Lasso) and L2 (Ridge) regularization were applied to enhance model stability and perform feature selection. To interpret the contribution of individual ROIs and TOIs to model predictions, we used SHAP (SHapley Additive exPlanations) values^59^. SHAP provides a game-theoretic framework to estimate the marginal impact of each feature on the predicted probability of affective blindsight, thereby identifying the most influential anatomical predictors^31,60^.

### Bayesian Statistical Modeling

To complement and validate these findings, we implemented a Bayesian logistic regression framework, which offers several advantages for small sample sizes and correlated predictors, including probabilistic inference, prior-based regularization, and robust model comparison via Bayes factors. All statistical analyses were performed in the R computing environment (version 4.3.2), utilizing the brms package for Bayesian regression modeling, bridgesampling for marginal likelihood estimation. Seven competing Bayesian generalized linear models were fitted to predict affective blindsight using a Bernoulli likelihood function with a logit link. These included: (1) a Null model with no lesion-defined tract predictors, (2) a Calcarine-only model (*Calcarine*) including only calcarine damage, (3) a Pulvinar–amygdala pathway model (*Pulvinar to Amygdala*) incorporating bilateral pulvinar–amygdala interactions, (4) a Pulvinar–pSTS pathway model (*Puvinar to pSTS*) comprising bilateral pulvinar–pSTS interactions, (5) a Left-Sided Pathways model (*Left sided*) with left pulvinar–amygdala and pulvinar–pSTS tract interactions, (6) a Right-Sided Pathways model (*Right sided*) with corresponding right hemisphere interactions, and (7) a full model (*Full Model*) incorporating calcarine damage along with all main effects and pairwise interactions among the four lesion-defined tracts. To ensure stable and interpretable parameter estimation, weakly informative priors were specified for all model components. The intercept was assigned a Student’s *t* distribution with 3 degrees of freedom, a mean of 0, and a scale parameter of 2.5, allowing for heavier tails to accommodate potential outliers. All tract-of-interest (TOI) predictors and the calcarine covariate were modeled using Normal priors with a mean of 0 and a standard deviation of 2.5, reflecting moderate prior uncertainty. Interaction terms were constrained using more regularizing Normal priors with a mean of 0 and a standard deviation of 1.5. Models were estimated using four MCMC chains of 8000 iterations each, with the first 2000 iterations per chain discarded as warm-up. A fixed random seed (1234) was used to ensure reproducibility. Sampler control parameters were adjusted to improve convergence and efficiency (adapt_delta = 0.95, max_treedepth = 15). Convergence was verified by inspecting R□ values (all < 1.01) and effective sample sizes. Marginal likelihoods were estimated for each model using the bridge_sampler method, and Bayes Factors (BFs) were computed relative to the Null Model. Posterior model probabilities, assuming equal prior model weights, were then derived from the BFs. A pairwise log□□ (Bayes Factor) comparison matrix was computed and visualized as a heatmap to illustrate the relative support for each model against every other. To validate the approach, we repeated the analysis using Bayesian logistic regression implemented through an in-house Python pipeline (see supp info).

### Lesion network mapping

Lesion network mapping is a validated technique that identifies regions functionally connected to a lesion location based on a fMRI resting state template. This allows one to localize symptoms even when lesions occur in various brain locations using the normative connectome data^61^. An open source normative resting state - fMRI dataset assembled from the 1000 healthy Brain Genomics Superstruct Project (https://dataverse.harvard.edu/dataverse/GSP) was used for lesion network mapping ^62^. Processing of the resting state fMRI data was performed with SPM-12 (Wellcome Department of Imaging Neuroscience, London, UK) and the CONN functional connectivity toolbox ^63^ both implemented in Matlab R2022a (The Math Works. Natick, MA, USA). The manually drawn lesion map was used as a seed in a resting state functional connectivity MRI analysis using the CONN toolbox. A connectivity r-map thus obtained for the lesion was converted to t-maps and thresholded at T > ±5 to create a lesion network map of significantly functionally connected regions to the patient’s lesion site (whole-brain voxel-wise FWE-corrected p < 0.05)^61^. The thresholded but non-binarized lesion network maps were used in a multivariate lesion network symptom mapping method using SVR LSM to identify the networks that were associated with the binary outcome of presence or absence of affective blindsight.

### Lesion disconnectome mapping

To study the impact of white matter tract disconnection, we performed disconnectome mapping to assess how lesions in one part of the cortex may affect distant regions through their structural connections. For each patient, disconnectome maps were generated using the BCBtoolkit, which leverages normative tractography data from healthy controls to estimate which white matter tracts are likely to be disrupted by a given lesion. This approach allowed us to infer the extent of structural disconnection beyond the lesion site itself and evaluate the involvement of specific tracts of interest ^64^. First, the lesion maps was normalised to MNI space and then used as a seed region for tractography to identify the white matter fibers that are anatomically connected to the lesion ^64^. The resulting disconnectome maps, which represent the probability of white matter disconnection for each voxel (ranging from 0% to 100%), were then incorporated into analyses of tracts of interest (TOIs) and used in multivariate lesion network-symptom mapping. Specifically, these maps were utilized in support vector regression–based lesion-symptom mapping (SVR-LSM) and in Bayesian lesion-deficit inference (BLDI) to examine the relationship between structural disconnection and affective blindsight. This procedure identified the damaged white matter pathways that were associated with the binary outcome of presence or absence of affective blindsight.

### Multivariate lesion–symptom mapping support vector regression

To identify neural correlates of affective blindsight, we performed support vector regression-based multivariate lesion–symptom mapping (SVR-LSM) using three types of imaging-derived maps: (1) binary lesion maps, (2) lesion network mapping, and (3) lesion disconnectome mapping, as described above. SVR-LSM was first implemented using the SVR-LSM toolbox (https://github.com/atdemarco/svrlsmgui) in MATLAB 2022a (The MathWorks Inc., Natick, MA), using a radial basis function kernel (gamma = 5, cost = 30)^65^. Voxels included in the analysis were restricted to those with lesion overlap in at least 10 patients. To validate and optimize the analysis, we repeated SVR-LSM using an in-house Python pipeline (https://github.com/cogneuro-rgb), performing a grid search with 5-fold cross-validation. The optimal hyperparameters identified were: epsilon = 0.1, cost = 10, gamma = 2. Lesion volume was regressed out of both the binary affective blindsight outcome and imaging maps. Lesion chronicity (acute vs. chronic) and other behavioral scores significantly associated with blindsight were included as covariates. Statistical significance was determined using 10,000 permutations with a voxel-wise threshold of p < 0.005 and cluster-level correction at p < 0.05. Identified clusters were anatomically localized using the Harvard–Oxford Cortical Atlas, the Automated Anatomical Labeling Atlas 3 (AAL3), and the Juelich Histological Atlas^66^.White matter tracts affected by lesions were identified using the IIT Human Brain Atlas.

### Bayesian Lesion–Deficit Inference (BLDI)

Given the relatively small sample size in our study of affective blindsight, we employed Bayesian Lesion–Deficit Inference (BLDI) to complement frequentist approaches^67^. Traditional frequentist methods can be limited in power when sample sizes are small or lesions are heterogeneous. BLDI offers a principled statistical framework better suited to low-powered settings, allowing for the detection of both positive and null lesion-deficit associations. We used the R package BLDI (https://github.com/ChrisSperber/BLDI) to perform voxel-wise Bayesian inference on three types of imaging-derived data: (1) binary lesion maps, (2) continuous lesion network connectivity maps, and (3) continuous disconnectome maps. BLDI models were implemented following established procedures and priors, with Bayes factors computed at each voxel to quantify the evidence for or against a lesion-deficit association^67^. The Bayesian framework has the added benefit of identifying regions where the data supports the absence of an association, information which is not available through frequentist inference alone^67^.

### Diffusion Tractography

A subset of three patients with affective blindsight who were available for a follow up visit, underwent diffusion tensor imaging (DTI). For processing the DTI data, we employed both global and local probabilistic tractography using MRtrix3 to reconstruct relevant white matter (WM) tracts ^68,69^. Preprocessing began with the extraction of the B0 image, which was coregistered to each patient’s T1-weighted anatomical image using FSL’s FLIRT^70^. The T1 image was then transformed into standard Montreal Neurological Institute (MNI) space, and this transformation was applied to the diffusion data to facilitate group-level analysis. Tissue segmentation was performed to generate five-tissue-type (5TT) images required for anatomically constrained tractography (ACT)^71^. Global tractography was first performed to model the whole-brain fiber architecture using a Bayesian generative approach. A tractogram of 50 million streamlines was generated and subsequently filtered to 10 million biologically meaningful streamlines using the Spherical-deconvolution Informed Filtering of Tractograms (SIFT) method. This filtering ensured that the streamline counts reflected the underlying fiber orientation distributions (FODs) and reduced the inherent bias toward longer or denser pathways. White matter FOD amplitudes were normalized across participants to allow for quantitative comparison of apparent fiber density (AFD) values. From the global tractogram, specific tracts were extracted by selecting streamlines that connected predefined bilateral regions of interest (ROIs) and excluding streamlines that passed through non-target ROIs. The ROIs included bilateral lateral geniculate nucleus (LGN), pulvinar, superior temporal sulcus (specifically the posterior superior temporal sulcus), human middle temporal visual area (hMT+), and the amygdala. The tracts that were studied included LGN–hMT+, pulvinar–hMT+, pulvinar–amygdala, and pulvinar-pSTStracts, reflecting pathways hypothesized to support non-conscious affective visual processing. Following global tractography, we conducted local probabilistic tractography using the iFOD2 algorithm. Here, 25,000 streamlines were seeded unidirectionally from each ROI to reconstruct connections in both directions (e.g., LGN→hMT+ and hMT+→LGN). Local tractography was confined to the relevant anatomical regions and employed ACT to improve biological plausibility by constraining streamlines to terminate at the gray matter–white matter interface. As in the global analysis, exclusion masks were used to ensure specificity of tracts—for example, excluding any LGN–hMT+ streamlines that traversed the pulvinar. To enhance the interpretability and reliability of tractography results, SIFT2 was applied during local tractography to assign cross-sectional multipliers to each streamline, providing a quantitative estimate of apparent fiber density that accounts for differences in tract volume and distribution. For visualization, individual tractography results were transformed into MNI space and combined for group-level display. Quantitative metrics—including streamline count, tract length, and normalized AFD—were extracted for each reconstructed tract and compared with seven age matched neurotypical control participants. This allowed us to evaluate whether the presence of affective blindsight was associated with preserved or altered structural connectivity within these subcortical and extrastriate visual pathways.

## Acknowledgments

We thank our patients for their participation, and Dr. Elise Rowe for proving tract masks for the ROI analysis.

## Conflict of Interest

The authors declare no competing financial or non-financial interests.

## Data Availability

De-identified data and analysis code are available https://osf.io/xue3d.

## Notes

https://osf.io/xue3d/

### Competing Interest Statement

The authors have declared no competing interest.

### Summary of Updates

This version of the manuscript has been revised to update the author names and details

## References

1. Weiskrantz, L., Warrington, E. K. K., Sanders, M. D. & Marshall, J. Visual capacity in the hemianopic field following a restricted occipital ablation. Brain 97, 709–728 (1974).

2. Weiskrantz, L. Blindsight: A Case Study and Its Implications. (Oxford University Press, Oxford, 1990).

3. Derrien, D., Garric, C., Sergent, C. & Chokron, S. The nature of blindsight: implications for current theories of consciousness. Neuroscience of Consciousness 2022, niab043 (2022).

4. Azzopardi, P. & Cowey, A. Blindsight and visual awareness. Conscious Cogn 7, 292–311 (1998).

5. Barbur, J. L., Weiskrantz, L. & Harlow, J. A. The unseen color aftereffect of an unseen stimulus: Insight from blindsight into mechanisms of color afterimages. Proceedings of the National Academy of Sciences 96, 11637–11641 (1999).

6. B, de G., J, V., G, P. & L, W. Non-conscious recognition of affect in the absence of striate cortex. Neuroreport 10, (1999).

7. Burra, N., Hervais-Adelman, A., Celeghin, A., de Gelder, B. & Pegna, A. J. Affective blindsight relies on low spatial frequencies. Neuropsychologia 128, 44–49 (2019).

8. Heywood, C. A. & Kentridge, R. W. Affective blindsight? Trends in Cognitive Sciences 4, 125–126 (2000).

9. Celeghin, A., de Gelder, B. & Tamietto, M. From affective blindsight to emotional consciousness. Consciousness and Cognition 36, 414–425 (2015).

10. Pegna, A. J., Khateb, A., Lazeyras, F. & Seghier, M. L. Discriminating emotional faces without primary visual cortices involves the right amygdala. Nat Neurosci 8, 24–25 (2005).

11. Morris, J. S., DeGelder, B., Weiskrantz, L. & Dolan, R. J. Differential extrageniculostriate and amygdala responses to presentation of emotional faces in a cortically blind field. Brain 124, 1241–1252 (2001).

12. Tamietto, M. & de Gelder, B. Neural bases of the non-conscious perception of emotional signals. Nat Rev Neurosci 11, 697–709 (2010).

13. de Gelder, B., van Honk, J. & Tamietto, M. Emotion in the brain: of low roads, high roads and roads less travelled. Nat Rev Neurosci 12, 425; author reply 425 (2011).

14. Tamietto, M. & de Gelder, B. Affective blindsight in the intact brain: Neural interhemispheric summation for unseen fearful expressions. Neuropsychologia 46, 820–828 (2008).

15. Rafal, R. D. et al. Connectivity between the superior colliculus and the amygdala in humans and macaque monkeys: virtual dissection with probabilistic DTI tractography. J Neurophysiol 114, 1947–1962 (2015).

16. Ajina, S., Pollard, M. & Bridge, H. The Superior Colliculus and Amygdala Support Evaluation of Face Trait in Blindsight. Frontiers in Neurology 11, (2020).

17. Pessoa, L. & Adolphs, R. Emotion and the brain: multiple roads are better than one. Nat Rev Neurosci 12, 425–425 (2011).

18. Seltzer, B. & Pandya, D. N. Afferent cortical connections and architectonics of the superior temporal sulcus and surrounding cortex in the rhesus monkey. Brain Research 149, 1–24 (1978).

19. Schobert, A.-K., Corradi-Dell’Acqua, C., Frühholz, S., van der Zwaag, W. & Vuilleumier, P. Functional organization of face processing in the human superior temporal sulcus: a 7T high-resolution fMRI study. Social Cognitive and Affective Neuroscience 13, 102–113 (2018).

20. Hein, G. & Knight, R. T. Superior Temporal Sulcus—It’s My Area: Or Is It? Journal of Cognitive Neuroscience 20, 2125–2136 (2008).

21. Pitcher, D. et al. The Human Posterior Superior Temporal Sulcus Samples Visual Space Differently From Other Face-Selective Regions. Cerebral Cortex 30, 778–785 (2020).

22. Pitcher, D. & Ungerleider, L. G. Evidence for a Third Visual Pathway Specialized for Social Perception. Trends Cogn Sci 25, 100–110 (2021).

23. Puce, A. From Motion to Emotion: Visual Pathways and Potential Interconnections. Journal of Cognitive Neuroscience 1–24 (2024) doi:10.1162/jocn_a_02141.

24. Prabhakar, A. T. et al. Double dissociation of dynamic and static face perception provides causal evidence for a third visual pathway. Nat Commun 16, 6513 (2025).

25. Iidaka, T., Miyakoshi, M., Harada, T. & Nakai, T. White matter connectivity between superior temporal sulcus and amygdala is associated with autistic trait in healthy humans. Neuroscience Letters 510, 154–158 (2012).

26. Aggleton, J. P., Burton, M. J. & Passingham, R. E. Cortical and subcortical afferents to the amygdala of the rhesus monkey (Macaca mulatta). Brain Res 190, 347–368 (1980).

27. Aggleton, J. P. & Mishkin, M. Visual impairments in macaques following inferior temporal lesions are exacerbated selectively by additional damage to superior temporal sulcus. Behavioural Brain Research 39, 262–274 (1990).

28. Andino, S. L. G., Menendez, R. G. de P., Khateb, A., Landis, T. & Pegna, A. J. Electrophysiological correlates of affective blindsight. Neuroimage 44, 581–589 (2009).

29. Striemer, C. L., Whitwell, R. L. & Goodale, M. A. Affective blindsight in the absence of input from face processing regions in occipital-temporal cortex. Neuropsychologia 128, 50–57 (2019).

30. McFadyen, J., Mattingley, J. B. & Garrido, M. I. An afferent white matter pathway from the pulvinar to the amygdala facilitates fear recognition. Elife 8, e40766 (2019).

31. Nohara, Y., Matsumoto, K., Soejima, H. & Nakashima, N. Explanation of machine learning models using shapley additive explanation and application for real data in hospital. Computer Methods and Programs in Biomedicine 214, 106584 (2022).

32. Kinoshita, M. et al. Dissecting the circuit for blindsight to reveal the critical role of pulvinar and superior colliculus. Nat Commun 10, 135 (2019).

33. de Gelder, B., Humphrey, N. & Pegna, A. J. On the bright side of blindsight. Considerations from new observations of awareness in a blindsight patient. Cerebral Cortex 35, 42–48 (2025).

34. Takakuwa, N., Isa, K., Onoe, H., Takahashi, J. & Isa, T. Contribution of the Pulvinar and Lateral Geniculate Nucleus to the Control of Visually Guided Saccades in Blindsight Monkeys. J Neurosci 41, 1755–1768 (2021).

35. Gerbella, M., Caruana, F. & Rizzolatti, G. Pathways for smiling, disgust and fear recognition in blindsight patients. Neuropsychologia 128, 6–13 (2019).

36. Bertini, C., Pietrelli, M., Braghittoni, D. & Làdavas, E. Pulvinar Lesions Disrupt Fear-Related Implicit Visual Processing in Hemianopic Patients. Frontiers in Psychology 9, (2018).

37. Gonzalez Andino, S. L., Grave de Peralta Menendez, R., Khateb, A., Landis, T. & Pegna, J. Electrophysiological correlates of affective blindsight. NeuroImage 44, 581–589 (2009).

38. Striemer, C. L., Whitwell, R. L. & Goodale, M. A. Affective blindsight in the absence of input from face processing regions in occipital-temporal cortex. Neuropsychologia 128, 50–57 (2019).

39. Kinoshita, M. et al. Dissecting the circuit for blindsight to reveal the critical role of pulvinar and superior colliculus. Nat Commun 10, 135 (2019).

40. Aleci, C. & Dutto, K. Seeing the invisible: theory and evidence of blindsight. Discov Med 1, 149 (2024).

41. Maldonado, I. L. et al. Multimodal study of multilevel pulvino-temporal connections: a new piece in the puzzle of lexical retrieval networks. Brain 147, 2245–2257 (2024).

42. Weiner, K. S. & Gomez, J. Third Visual Pathway, Anatomy, and Cognition across Species. Trends in Cognitive Sciences 25, 548–549 (2021).

43. Pitcher, D. Neuropsychological evidence of a third visual pathway specialized for social perception. Nat Commun 16, 5774 (2025).

44. Pitcher, D., Duchaine, B. & Walsh, V. Combined TMS and fMRI Reveal Dissociable Cortical Pathways for Dynamic and Static Face Perception. Current Biology 24, 2066–2070 (2014).

45. Elorette, C., Forcelli, P. A., Saunders, R. C. & Malkova, L. Colocalization of Tectal Inputs With Amygdala-Projecting Neurons in the Macaque Pulvinar. Front Neural Circuits 12, 91 (2018).

46. Kaas, J. H. & Lyon, D. C. Pulvinar Contributions to the Dorsal and Ventral Streams of Visual Processing in Primates. Brain Res Rev 55, 285–296 (2007).

47. Kohler, C. G. et al. Facial emotion recognition in schizophrenia: intensity effects and error pattern. Am J Psychiatry 160, 1768–1774 (2003).

48. Gilbert, M., Demarchi, S. & Urdapilleta, I. FACSHuman, a software program for creating experimental material by modeling 3D facial expressions. Behavior Research Methods (2021) doi:10.3758/s13428-021-01559-9.

49. Vaina, L. M. et al. Functional and anatomical profile of visual motion impairments in stroke patients correlate with fMRI in normal subjects. J Neuropsychol 4, 121–145 (2010).

50. Siuda-Krzywicka, K. et al. Color Categorization Independent of Color Naming. Cell Reports 28, 2471-2479.e5 (2019).

51. Newcombe, F., Oldfield, R. C., Ratcliff, G. G. & Wingfield, A. Recognition and naming of object-drawings by men with focal brain wounds. J Neurol Neurosurg Psychiatry 34, 329–340 (1971).

52. Valentine, T. & Bruce, V. Mental rotation of faces. Memory & Cognition 16, 556–566 (1988).

53. rordenlab/MRIcroGL. Chris Rorden’s Lab (2025).

54. FreeSurferVersion3 - Free Surfer Wiki. https://surfer.nmr.mgh.harvard.edu/fswiki/FreeSurferVersion3.

55. Iglesias, J. E. et al. A probabilistic atlas of the human thalamic nuclei combining ex vivo MRI and histology. NeuroImage 183, 314–326 (2018).

56. Desikan, R. S. et al. An automated labeling system for subdividing the human cerebral cortex on MRI scans into gyral based regions of interest. NeuroImage 31, 968–980 (2006).

57. Wang, L., Mruczek, R. E. B., Arcaro, M. J. & Kastner, S. Probabilistic Maps of Visual Topography in Human Cortex. Cereb Cortex 25, 3911–3931 (2015).

58. Rowe, E. G., Zhang, Y. & Garrido, M. I. Evidence for adaptive myelination of subcortical shortcuts for visual motion perception in healthy adults. Hum Brain Mapp 44, 5641–5654 (2023).

59. Lundberg, S. et al. atprabhakar/shap: Shap. Zenodo 10.5281/zenodo.17341203 (2025).

60. Ponce‐Bobadilla, A. V., Schmitt, V., Maier, C. S., Mensing, S. & Stodtmann, S. Practical guide to SHAP analysis: Explaining supervised machine learning model predictions in drug development. Clin Transl Sci 17, e70056 (2024).

61. Boes, A. D. et al. Network localization of neurological symptoms from focal brain lesions. Brain 138, 3061–3075 (2015).

62. Buckner, R. L., Roffman, J. L. & Smoller, J. W. Brain Genomics Superstruct Project (GSP). Harvard Dataverse 10.7910/DVN/25833 (2020).

63. Whitfield-Gabrieli, S. & Nieto-Castanon, A. Conn: A Functional Connectivity Toolbox for Correlated and Anticorrelated Brain Networks. Brain Connectivity 2, 125–141 (2012).

64. Foulon, C. et al. Advanced lesion symptom mapping analyses and implementation as BCBtoolkit. Gigascience 7, (2018).

65. DeMarco, A. T. & Turkeltaub, P. E. A multivariate lesion symptom mapping toolbox and examination of lesion-volume biases and correction methods in lesion-symptom mapping. Human Brain Mapping 39, 4169–4182 (2018).

66. Amunts, K., Mohlberg, H., Bludau, S. & Zilles, K. Julich-Brain: A 3D probabilistic atlas of the human brain’s cytoarchitecture. Science 369, 988–992 (2020).

67. Sperber, C., Gallucci, L., Smaczny, S. & Umarova, R. Bayesian lesion-deficit inference with Bayes factor mapping: key advantages, limitations, and a toolbox. Neuroimage 120008 (2023) doi:10.1016/j.neuroimage.2023.120008.

68. MRtrix3. https://www.mrtrix.org/.

69. Tournier, J.-D. et al. MRtrix3: A fast, flexible and open software framework for medical image processing and visualisation. NeuroImage 202, 116137 (2019).

70. Jenkinson, M., Bannister, P., Brady, M. & Smith, S. Improved optimization for the robust and accurate linear registration and motion correction of brain images. Neuroimage 17, 825–841 (2002).

71. Smith, R. E., Tournier, J.-D., Calamante, F. & Connelly, A. Anatomically-constrained tractography: Improved diffusion MRI streamlines tractography through effective use of anatomical information. NeuroImage 62, 1924–1938 (2012).

