## Supplementary material for "Multimodal lesion mapping in affective blindsight reveals dual amygdala and superior temporal sulcus contributions to nonconscious emotion processing": Supp Info

**The PDF file includes:**

Supplementary Text

Figs. S1 to S3

Tables S1 to S4

**Figure S.1. Regions of Interest(ROI) Masks used in the analysis**

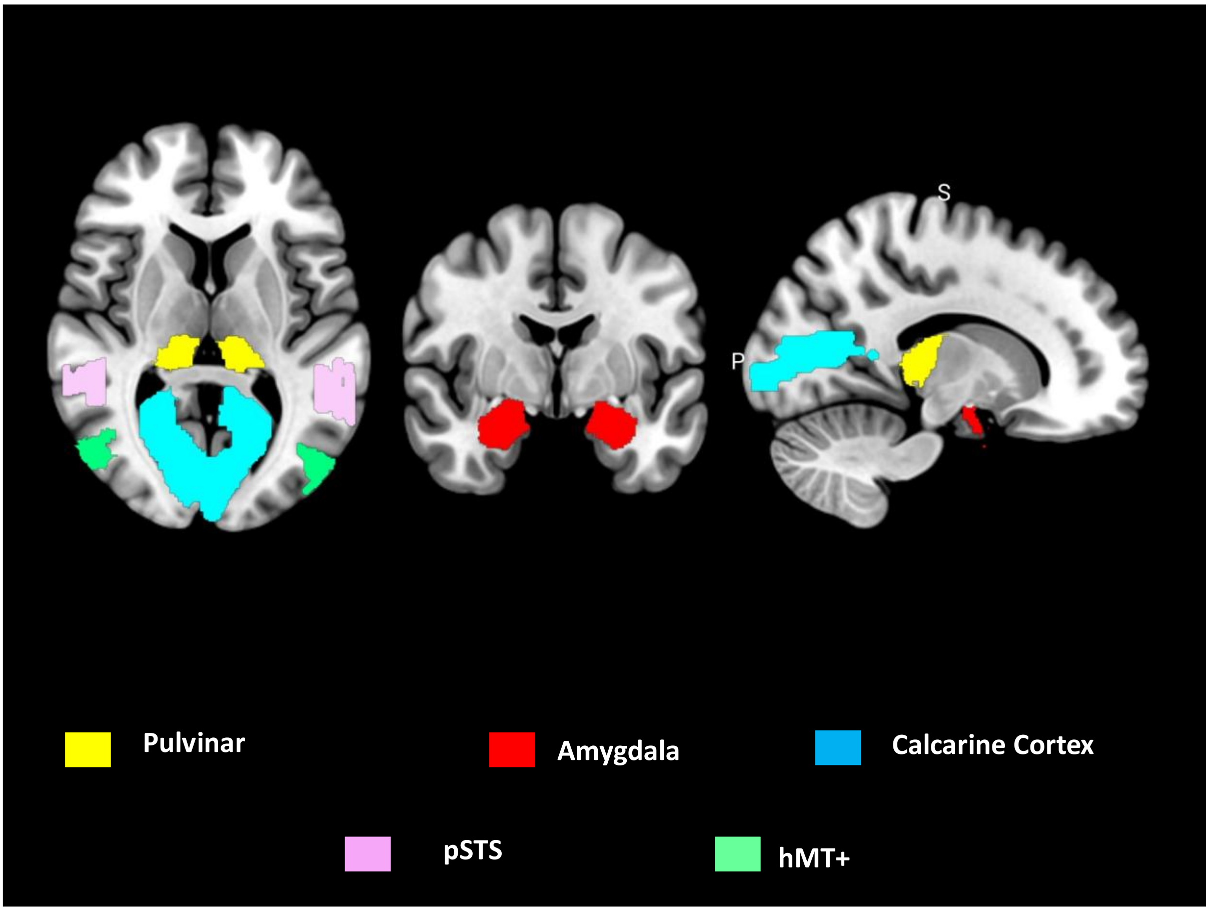


Figure S.1. Shows the Regions of Interest (ROIs) Used in Lesion Analysis plotted on the MNI152 template brain. **hMT+**- human motion-selective area, **pSTS** - posterior superior temporal sulcus (pSTS)

### Behavioral Results

A total of 182 patients with higher-order visual deficits following focal brain lesions were screened for inclusion. Of these, 31 patients were identified to have bilateral lesions to the calcarine cortex were identified and included in the analysis (Fig.S2).

**Figure S2– Flow Diagram of recruited patients**

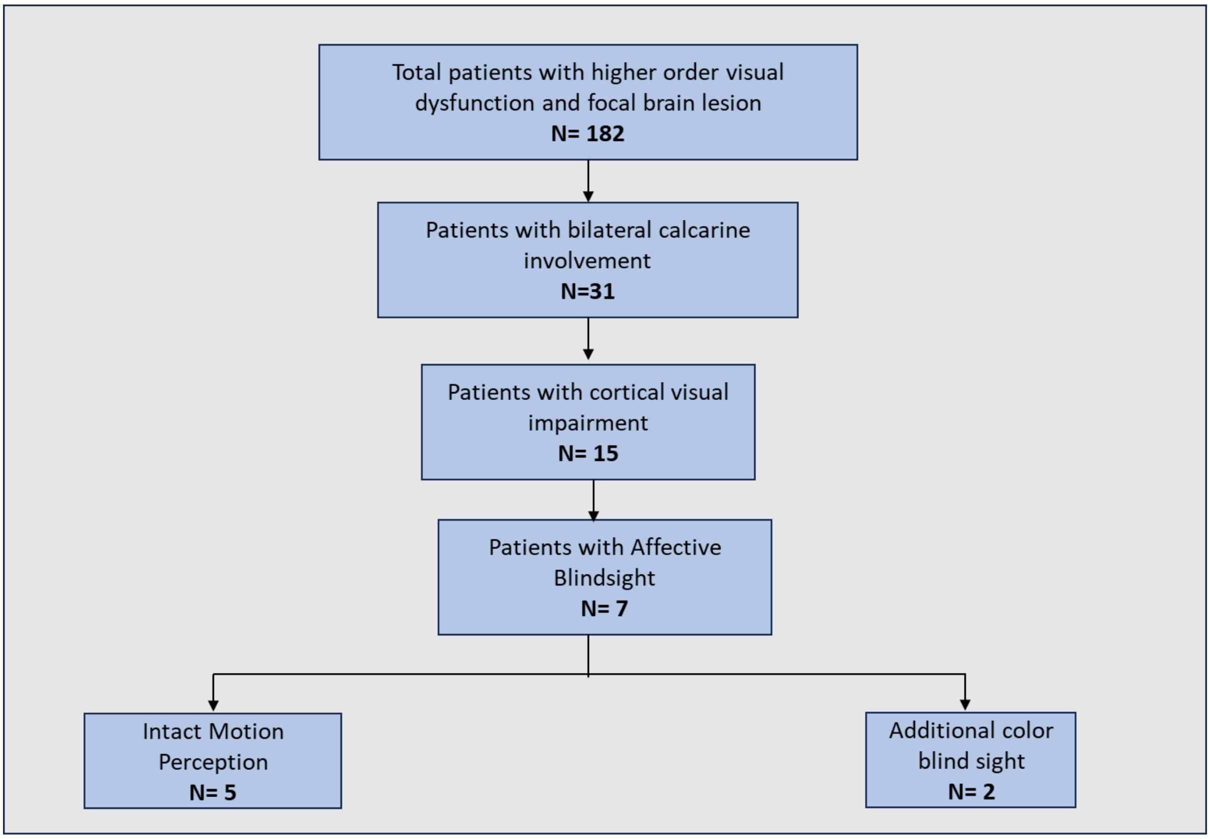


Flow diagram illustrates the systematic screening and selection of patients with affective blindsight from an initial cohort with higher-order visual dysfunction and focal brain lesions.

Among these, 29 had ischemic strokes confined to the posterior cerebral artery (PCA) territory and 2 had a chronic unspecified posterior leukoencephalopathy. Of these, 3 patients had acute lesions (<14 days post-ictus), and 28 patients had chronic lesions (>6 months post-ictus). In several cases (Patients 1, 3, and 4), patients reported intermittent awareness of visual stimuli—such as red-colored objects or faces—described as brief percepts against a hazy or blank visual background that disappeared within seconds. Most patients (Patients 2–6) performed significantly above chance on dynamic facial emotion recognition tasks using a two-alternative forced choice (2AFC) design, despite denying any visual awareness during testing. One patient (Patient 3) initially presented with Anton syndrome, showing visual anosognosia despite complete cortical blindness, which later transitioned to demonstrable blindsight for both motion and affect. Longitudinal follow-up assessments (Patients 3 and 6) revealed partial recovery of some visual functions, particularly for motion and coarse shape outlines, while deficits in color vision and face identification persisted.

**Figure S3. Lesion coverage displayed in the MNI template Brain**

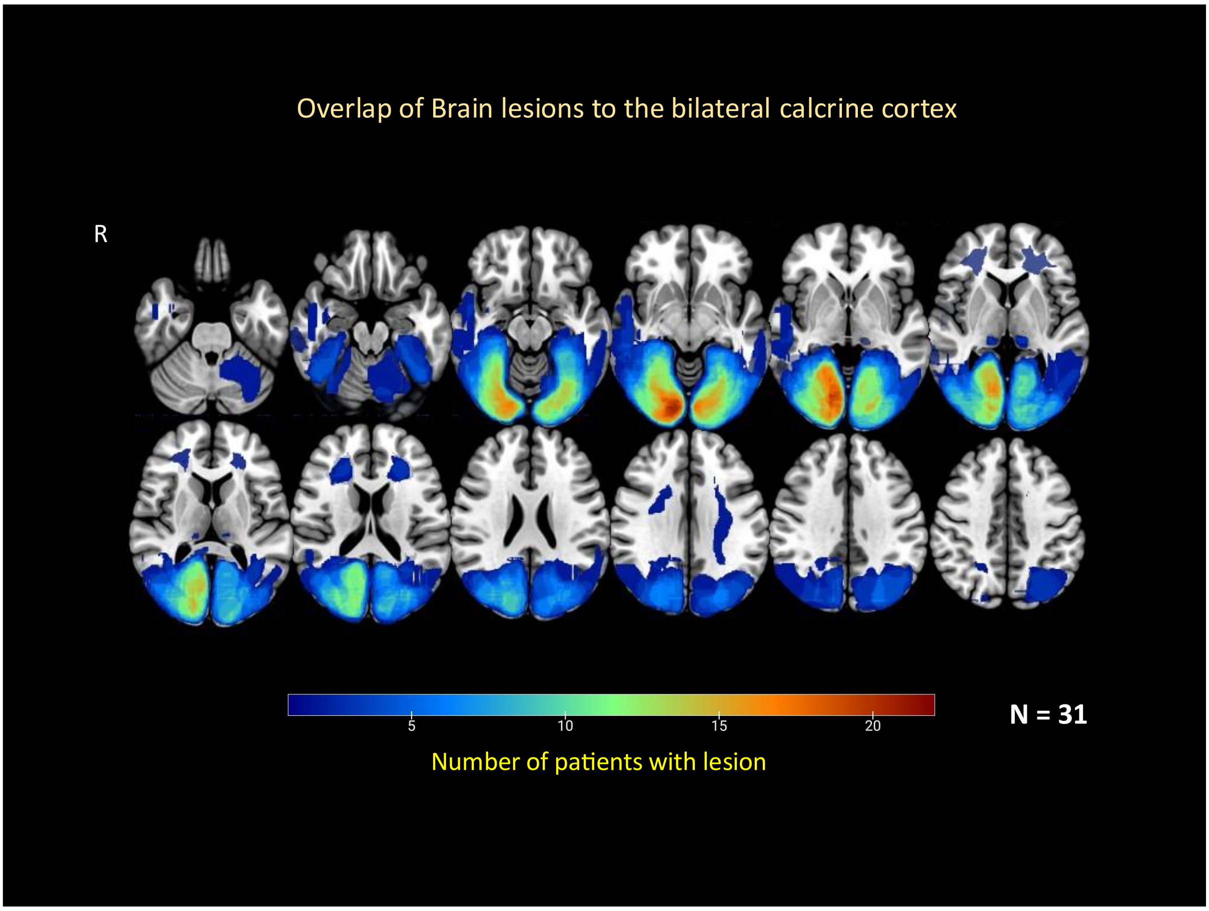


Lesion overlay map for 31 patients with bilateral calcarine lesions. The color-coded legend denotes the number of patients with damage to a specific brain region. The lesion maps are displayed on the axial slices of the MNI 152 template brain oriented in the neurological convention. R- denotes the right side of the brain images.

### Clinical Profiles and Lived Experience of Patients With and Without Affective Blindsight

We present detailed clinical profiles and lived experiences of 15 patients with profound cortical visual impairment due to acquired brain injuries. Patients 1–7 demonstrated preserved non-conscious affective processing consistent with affective blindsight, while Patients 8–15 exhibited severe visual deficits without evidence of preserved non-conscious emotional processing. All patients underwent comprehensive neuro-ophthalmological evaluation, neuroimaging, and standardized behavioral testing using two-alternative forced choice paradigms wherever feasible.

**Patients With Affective Blindsight (Patients 1–7)**

**Patient 1** was a 58-year-old male farmer who presented to our center approximately one year following the onset of acute visual loss. He had a prior history of a left frontoparietal infarct in the middle cerebral artery (MCA) territory 1.5 years earlier, which had resulted in persistent right-sided spastic hemiparesis. Approximately six months after the first event, he experienced a second event. He reported awakening from sleep with sudden-onset complete visual loss, which was preceded by brief, ill-formed episodes of flashing white lights in his visual field. Since then, he described being functionally blind under normal viewing conditions. Occasionally, he reported brief episodes of awareness of visual objects—often red-coloured items or faces—that would appear against a diffuse white haze and vanish within seconds. Although he could detect their presence, he was unable to recognize the faces or objects and could not determine their spatial location. Ophthalmologic examination revealed no ocular pathology: visual acuity could not be reliably assessed due to cortical impairment; pupillary responses were intact with no relative afferent pupillary defect (RAPD); and fundus examination was normal. He was unable to recognize or match common objects or colors and failed on standard tests of visual naming. Despite this, motion perception was relatively preserved, as evidenced by normal performance on dot motion direction discrimination. On testing for emotion recognition, he denied any visual awareness during task performance, but he was able to identify basic emotional expressions on the 2AFC tasks. Similarly, he performed above chance in color discrimination tasks.

**Patient 2** was a 72-year-old right-handed male who was evaluated two years following a posterior circulation ischemic stroke secondary to basilar artery stenosis in the context of intracranial and extracranial atherosclerotic disease. He was a poorly controlled type 2 diabetic with systemic hypertension. The index event was characterized by acute-onset vertigo, visual disturbances, and ataxia, consistent with brainstem and occipital involvement. Initial neuroimaging confirmed bilateral occipital infarcts and cerebellar involvement. Over the following months, the patient reported persistent visual impairment that failed to improve substantially. At the time of evaluation, he had a vague sense of light and motion but was not able to see form or color. Examination revealed preserved pupillary responses and normal ocular fundus. Behavioral testing using two-alternative forced-choice (2AFC) paradigms revealed preserved non-conscious affective processing.

**Patient 3** was a 46-year-old right-handed male who presented with an acute posterior circulation infarction consistent with top-of-the-basilar artery syndrome. His clinical presentation included abrupt onset of visual loss and altered awareness. Neuroimaging revealed bilateral occipital lobe infarcts, with associated involvement of the bilateral thalami, left medial temporal lobe, and the left posterior capsuloganglionic region. Vascular imaging demonstrated occlusion of the left posterior cerebral artery (P2–P4 segments) and critically reduced flow in the distal branches of the right posterior cerebral artery(PCA). The underlying etiology was suspected to be either cardioembolic or secondary to radiation-induced vasculopathy, in the context of prior neck radiotherapy for oral carcinoma. At initial presentation, the patient exhibited complete cortical blindness, accompanied by visual anosognosia. Despite being unable to detect light or motion, he clearly denied any visual impairment and consistently offered high-confidence but incorrect responses on visual tasks—a classic presentation of Anton syndrome. No ocular pathology was identified on examination, and pupillary light responses were preserved. Longitudinal behavioral assessments were systematically performed at multiple time points post-stroke: days 1, 7, 10, and 14; as well as at 1 month, 3 months, 1 year, and 2 years. On day 7, he remained cortically blind, without any demonstrable visual awareness, yet continued to exhibit anosognosia. By day 10, he began to perform above chance in direction-of-light detection tasks, suggesting the emergence of rudimentary non-conscious visual processing. By day 14, more extensive behavioral testing revealed the presence of residual unconscious visual abilities. He was able to discriminate the direction of coherent motion and identify affective expressions in dynamic facial stimuli at above-chance levels on two-alternative forced-choice (2AFC) paradigms, indicating the presence of both motion and affective blindsight. However, color discrimination and object recognition remained severely impaired. He was unable to name or match colors and failed to identify or differentiate faces, including an inability to distinguish normal from jumbled faces. At the three-month follow-up, the patient reported subjective improvement in visual awareness, describing intermittent perception of light and outlines. He also exhibited improvement in visually guided actions, such as reaching and grasping, suggestive of recovered visuomotor integration through dorsal stream pathways. While he reported perceiving the presence of faces, he remained unable to identify individuals, consistent with persistent prosopagnosia. Notably, performance in affective processing tasks using dynamic facial stimuli remained stable and above chance.

By one- and two-years post-stroke, gradual recovery of object recognition and naming abilities was observed, though significant deficits persisted. Achromatopsia remained prominent, and color perception did not recover. Prosopagnosia also remained unchanged; the patient continued to fail on face recognition tasks, both in naturalistic settings and formal testing. Motion perception and affective blindsight, however, remained preserved and stable over time. Diffusion MRI and tractography was performed at the two-year follow-up visit.

**Patient 4** was a 59-year-old right-handed male with a history of multiple cerebrovascular events, including a right hemispheric ischemic stroke followed by a posterior circulation infarct 7 years prior. He was evaluated seven years after the latter event for persistent and debilitating visual impairment. At the time of assessment, he reported near-complete visual loss. Subjectively, he described his visual experience as consisting predominantly of a uniform haze, with only occasional perception of ambient light or vague outlines of large objects. He denied any recognition of faces or objects and had minimal awareness of color or detailed visual form. However, he noted a subtle and intermittent sense of motion occurring within the environment, particularly during dynamic scenes. Neurological examination revealed residual right-sided spastic hemiparesis with impaired fine motor movements distally. Fundoscopy and pupillary reflexes were normal, and there was no relative afferent pupillary defect.

Formal behavioral testing revealed profound impairments in object and color recognition. On tasks involving color naming and matching, performance was at chance levels, consistent with achromatopsia. He was found to have intact motion discrimination and affective blindsight based on the 2AFC dynamic emotion recognition task.

**Patient 5** was a 63-year-old man who presented with a 15-year history of behavioral changes, memory disturbances, visual impairment, and gait abnormalities. Neuroimaging revealed extensive bilateral occipital white matter changes, more prominent on the left, consistent with chronic posterior leukoencephalopathy. At the time of evaluation, he reported complete visual loss, with no subjective awareness of form, light direction, motion, or facial outlines. On formal behavioral testing, he was unable to detect motion or distinguish between faces and objects, and could not differentiate normal from jumbled faces. Despite this, he demonstrated above-chance performance on a two-alternative forced choice (2AFC) dynamic emotion recognition task. Similarly, his performance on color naming tasks in a 2AFC format was above chance, indicating residual chromatic processing.

**Patient 6** was a 35-year-old man with a history of mitral valve replacement 12 years prior, who presented with acute onset headache followed by a generalized tonic-clonic seizure and visual disturbances. Neuroimaging revealed bilateral occipital lobe infarcts, and a cardioembolic etiology was suspected. In the acute post-stroke period, he exhibited complete cortical blindness, reporting no awareness of light, contours, motion, or spatial location. He was unable to navigate independently and required continuous assistance. On day 14 post-stroke, he reported vague light perception, although directionality and form remained absent. By the three-month follow-up, he described being able to perceive outlines and silhouettes of objects in well-lit environments. However, he remained unable to consciously recognize or name objects, faces, or colors. Subjective awareness of visual stimuli remained minimal, with a persistent sense of visual haze despite some functional improvement. Behavioral testing at 14 days and one-month post-stroke revealed preserved color perception and affective blindsight. Despite denying any conscious awareness of visual stimuli, he consistently performed above chance on a two-alternative forced choice (2AFC) paradigm involving dynamic facial expressions.

**Patient 7** was a 22-year-old woman with a history of deliberate self-harm ingested monocrotophos insecticide, resulting in cardiac arrest and subsequent hypoxic-ischemic encephalopathy with watershed infarcts. She presented three months later with severe visual impairment, describing perception of only a diffuse haze without recognition of objects, faces, or movement. MRI revealed bilateral occipitotemporal hyperintensities, affecting regions associated with both dorsal and ventral visual streams. On initial neuropsychological testing, she was unable to recognize objects, faces, or detect motion, and was impaired in static emotion recognition. Color perception was preserved. She showed above-chance performance on dynamic emotion recognition only during a two-alternative forced choice (2AFC) task, suggesting residual non-conscious affective processing. At follow-up six months later, despite persistent deficits in object recognition, motion perception, and visuomotor integration, she demonstrated spontaneous and accurate recognition of dynamic emotional expressions, even in the absence of intact motion perception.

**Patients Without Affective Blindsight (Patients 8–15)**

**Patient 8** was a 69-year-old man with a history of recurrent ischemic strokes involving the occipital lobes and type 2 diabetes mellitus. His first stroke occurred in 4 years ago and resulted in an altitudinal visual field defect and tunnel vision. Following a second occipital event one year later, he became completely blind, with only a vague and non-localizable perception of light. He reported no awareness of movement, form, or faces. Formal behavioral testing using two-alternative forced choice paradigms was attempted but not feasible due to his profound impairment.

**Patient 9** was a 52-year-old man who presented with sudden, painless bilateral visual loss accompanied by mild headache and giddiness. On admission, he had profound visual impairment and his ocular examination, including fundus assessment, was unremarkable. Magnetic resonance imaging of the brain revealed multiple infarcts in the posterior circulation. Evaluation identified poorly controlled diabetes (HbA1c 7.2%) and dyslipidemia as vascular risk factors. He remained unable to recognize form or color, though he reported some awareness of motion. At one-year follow-up, he continued to experience profound visual loss with minimal improvement, and formal behavioral testing was not possible.

**Patient 10** was a 54-year-old former military serviceman who presented with a sudden onset of vertiginous sensation followed by progressive impairment of consciousness over the course of one day. He was last seen well in the morning, but by evening was found unresponsive and brought to the emergency department. Neuroimaging revealed extensive bilateral occipital involvement. At two weeks post-onset, he remained unable to perceive light, form, motion, or color. He was unable to engage in behavioral testing due to the severity of his deficits. At one-and-a-half years of follow-up, there was no improvement in visual function, and he continued to report complete blindness.

**Patient 11** was a 52-year-old man who experienced a severe posterior circulation ischemic stroke, resulting in bilateral occipital infarcts and persistent cortical blindness. At the time of his initial assessment, he reported a complete absence of visual perception. Despite rehabilitation and follow-up evaluations, his visual status remained unchanged, and he was unable to participate in two-alternative forced choice testing.

**Patient 12** was a 69-year-old man with longstanding diabetes and systemic hypertension who developed severe cortical visual loss following recurrent posterior circulation strokes. He reported an initial period of tunnel vision after the first stroke, which progressed to complete blindness after a second event. He described a vague awareness of light but no perception of form, color, or motion. Despite some subjective improvement over several years, his deficits remained profound, and his performance was not above chance in any of the 2AFC tasks.

**Patient 13** was a 52-year-old man who presented with acute bilateral visual loss, accompanied by vertigo and headache. Initial examination revealed visual acuity of 6/24 with a normal fundus appearance. Brain imaging demonstrated multiple posterior circulation infarcts. Despite treatment, his vision rapidly worsened, and he developed tunnel vision followed by complete visual loss by the second day of admission. At one year, he was unable to identify form or color but could occasionally detect motion. Behavioral Performance not above chance in any of the 2AFC tasks

**Patient 14** was a 54-year-old man, a former military serviceman, who experienced an acute episode of vertigo followed by a prolonged period of unresponsiveness. He was found unconscious at his workplace and brought to the hospital. Imaging confirmed extensive occipital involvement. Two weeks after symptom onset, he was unable to perceive light, form, color, or movement. At one-and-a-half years of follow-up, his visual deficits remained unchanged, with no signs of recovery. He was unable to complete the 2AFC tasks.

**Patient 15** was a 69-year-old man with a history of recurrent posterior circulation strokes and poorly controlled diabetes. Following his second occipital infarct, he became completely blind. He reported only a vague sense of illumination, without the ability to localize light or detect movement. Despite subjective reports of partial improvement over time, his visual impairment remained profound, and he was unable to complete the 2AFC tasks.

### Diffusion tractography results

Tractography analysis was conducted in three patients (Patients 1, 2, and 3) with affective blindsight to assess the integrity of key subcortical–cortical visual pathways. As shown in the tables (see below Table S1 to S4), there was a general reduction in fiber count and Apparent Fiber Density (AFD) in the patient group compared to matched controls. Nonetheless, the presence of tracts connecting the pulvinar to the pSTS and amygdala in these patients supports the potential role of these pathways in supporting residual affective processing despite primary visual cortex damage

**Table S 1 Results of the Negative Binomial Regression – comparing the Global Tractography streamline counts between three patients with blindsight and neurotypical controls**

| Tract | Control Mean ± SD | Patient Mean ± SD | IRR (95% CI) | p-value |
|---|---|---|---|---|
| **Left LGN → Left MT** | 6 ± 8.12 | 1 ± 1.73 | 0.17 (0.02 – 1.94) | 0.126 |
| **Left pSTS → Left MT** | 29.5 ± 29.87 | 25.67 ± 11.68 | 0.87 (0.26 – 3.14) | 0.823 |
| **Left pulvinar → Left amygdala** | 814.25 ± 1144.66 | 617.33 ± 675.50 | 0.76 (0.10 – 7.11) | 0.783 |
| **Left pulvinar → Left MT** | 31.5 ± 47.28 | 1.67 ± 2.08 | 0.05 (0.02 – 0.12) | <0.0001 *** |
| **Left pulvinar → Left pSTS** | 9.25 ± 4.35 | 0.67 ± 1.15 | 0.07 (0.01 – 0.26) | 0.0006 *** |
| **Right LGN → Right MT** | 41 ± 48.99 | 1 ± 1.73 | 0.02 (0.001 – 0.76) | 0.0127 * |
| **Right pSTS → Right MT** | 153 ± 204.51 | 96 ± 70.41 | 0.63 (0.12 – 3.80) | 0.578 |
| **Right pulvinar → Right amygdala** | 1339.5 ± 1523.43 | 1283 ± 2173.04 | 0.96 (0.92 – 0.99) | 0.0414 * |
| **Right pulvinar → Right MT** | 50.5 ± 71.06 | 3.67 ± 2.52 | 0.07 (0.01 – 0.48) | 0.0035 ** |
| **Right pulvinar → Right pSTS** | 71 ± 50.88 | 6 ± 5.20 | 0.08 (0.05 – 0.13) | <0.0001 *** |

### Table S2. Results of the Negative Binomial Regression – comparing the Local Tractography streamline counts between three patients with blindsight and neurotypical controls

| Tract | Control Mean ± SD | Patient Mean ± SD | IRR (95% CI) | p-value |
|---|---|---|---|---|
| **Left LGN → Left MT** | 3114 ± 5734.85 | 1808.33 ± 2801.65 | 0.58 (0.05 – 8.82) | 0.645 |
| **Left pSTS → Left MT** | 3752.5 ± 2744.07 | 3787 ± 1489.14 | 1.01 (0.27 – 4.09) | 0.989 |
| **Left pulvinar → Left amygdala** | 14576 ± 7300.28 | 18609.33 ± 2557.73 | 1.28 (0.68 – 2.43) | 0.447 |
| **Left pulvinar → Left MT** | 3397.5 ± 5914.43 | 618.33 ± 536.02 | 0.18 (0.02 – 2.05) | 0.113 |
| **Left pulvinar → Left pSTS** | 1266 ± 1388.32 | 284.67 ± 305.97 | 0.22 (0.05 – 1.14) | 0.051 |
| **Right LGN → Right MT** | 7502.5 ± 7132.10 | 349.33 ± 387.04 | 0.05 (0.01 – 0.38) | 0.001 |
| **Right pSTS → Right MT** | 11956.25 ± 7982.61 | 7377 ± 8169.87 | 0.62 (0.15 – 2.76) | 0.497 |
| **Right pulvinar → Right amygdala** | 13051.5 ± 9963.46 | 10259 ± 10309.85 | 0.79 (0.78 – 0.80) | <0.0001 |
| **Right pulvinar → Right MT** | 5540 ± 8853.44 | 429.33 ± 505.34 | 0.08 (0.01 – 0.56) | 0.005 |
| **Right pulvinar → Right pSTS** | 13476.75 ± 9020.24 | 1482.33 ± 1304.65 | 0.11 (0.11 – 0.11) | <0.0001 |

### Table S3. Results of the Beta Binomial Regression – comparing the Global Tractography Apparent fiber density(AFD) between three patients with blindsight and neurotypical controls

| Tract | Control Mean ± SD | Patient Mean ± SD | Beta (95% CI) | p-value |
|---|---|---|---|---|
| **Left LGN → Left MT** | 0.48 ± 0.36 | 0.33 ± 0.58 | −0.20 (−2.24 – 1.84) | 0.847 |
| **Left pSTS → Left MT** | 0.61 ± 0.02 | 0.61 ± 0.13 | 0.04 (−0.40 – 0.48) | 0.870 |
| **Left pulvinar → Left amygdala** | 0.51 ± 0.07 | 0.52 ± 0.07 | 0.02 (−0.32 – 0.36) | 0.904 |
| **Left pulvinar → Left MT** | 0.29 ± 0.33 | 0.53 ± 0.49 | 0.53 (−1.37 – 2.44) | 0.583 |
| **Left pulvinar → Left pSTS** | 0.70 ± 0.09 | 0.21 ± 0.36 | −3.61 (−5.59 – −1.63) | **0.0004** |
| **Right LGN → Right MT** | 0.55 ± 0.20 | 0.28 ± 0.32 | −1.78 (−3.30 – −0.26) | **0.023** |
| **Right pSTS → Right MT** | 0.67 ± 0.03 | 0.65 ± 0.08 | −0.26 (−0.65 – 0.14) | 0.194 |
| **Right pulvinar → Right amygdala** | 0.51 ± 0.07 | 0.52 ± 0.07 | 0.10 (−0.23 – 0.43) | 0.531 |
| **Right pulvinar → Right MT** | 0.20 ± 0.26 | 0.43 ± 0.42 | 0.56 (−1.11 – 2.23) | 0.496 |
| **Right pulvinar → Right pSTS** | 0.64 ± 0.09 | 0.22 ± 0.34 | −2.64 (−4.47 – −0.82) | **0.0048** |

**Table S 4. Results of the Beta Binomial Regression – comparing the Local Tractography Apparent fiber density(AFD) between three patients with blindsight and neurotypical controls**

| Tract | Control Mean ± SD | Patient Mean ± SD | Beta (95% CI) | p-value |
|---|---|---|---|---|
| **Left LGN → Left MT** | 0.67 ± 0.13 | 0.80 ± 0.06 | 0.64 (−0.04 – 1.33) | 0.066 |
| **Left pSTS → Left MT** | 0.62 ± 0.02 | 0.61 ± 0.06 | −0.06 (−0.28 – 0.15) | 0.562 |
| **Left pulvinar → Left amygdala** | 0.51 ± 0.06 | 0.52 ± 0.08 | 0.04 (−0.31 – 0.38) | 0.833 |
| **Left pulvinar → Left MT** | 0.64 ± 0.03 | 0.73 ± 0.11 | 0.47 (−0.01 – 0.96) | 0.057 |
| **Left pulvinar → Left pSTS** | 0.72 ± 0.11 | 0.63 ± 0.04 | −0.45 (−0.997 – 0.097) | 0.107 |
| **Right LGN → Right MT** | 0.62 ± 0.13 | 0.71 ± 0.09 | 0.41 (−0.25 – 1.07) | 0.219 |
| **Right pSTS → Right MT** | 0.60 ± 0.07 | 0.52 ± 0.09 | −0.32 (−0.73 – 0.09) | 0.127 |
| **Right pulvinar → Right amygdala** | 0.51 ± 0.08 | 0.56 ± 0.10 | 0.20 (−0.24 – 0.64) | 0.369 |
| **Right pulvinar → Right MT** | 0.64 ± 0.11 | 0.59 ± 0.04 | −0.23 (−0.72 – 0.27) | 0.373 |
| **Right pulvinar → Right pSTS** | 0.60 ± 0.06 | 0.62 ± 0.08 | 0.09 (−0.28 – 0.46) | 0.635 |

### Comparison with Null tractography

As tractography is inferential in nature, we sought to objectively evaluate each hypothesized pathway by comparing targeted tractography outcomes with a corresponding null distribution. Null tractography was implemented by generating 100,000 streamlines from each ROI without incorporating fibre orientation information, while maintaining the same anatomical constraints as conventional tractography, yielding up to 200,000 streamlines per tract as a null benchmark (Table S5). To formally compare streamline counts from FOD-based and null tractography, we employed negative binomial regression, including Euclidean distance between ROIs as a covariate. This analysis revealed no evidence that the pulvinar–pSTS pathway could be reconstructed beyond anatomical proximity; it appeared stronger in null tractograms, likely reflecting the close spatial relationship of the ROIs. By contrast, FOD-based tractography, which enforces stricter anatomical constraints, produced fewer streamlines for closely adjacent regions. While tractography remains probabilistic, cadaveric dissection provides the reference standard for white matter pathways. Importantly, cadaveric studies have confirmed the pulvinar–pSTS pathway, and in all three patients who underwent DTI, both T1- and T2-weighted MRI consistently demonstrated intact white matter in this region[41]. Taken together, these findings suggest that our demonstration of pulvinar–pSTS tracts reflects preserved rather than artifactual connections.

**Table S5: Results of the Negative Binomial Regression – comparing the Data driven Tractography with Null Tractography**

| **Local Tractography** | | | | |
|---|---|---|---|---|
| **Tract** | **Actual Mean ± SD** | **Null Mean ± SD** | **IRR (95% CI)** | **p-value** |
| **Left pulvinar → Left amygdala** | 22621±434.14 | 7994±13089.02 | 2.83 (0.29 – 27.99) | 0.330 |
| **Left pulvinar → Left pSTS** | 346.33±436.02 | 1913±1901.41 | 0.18 (0.04 – 0.92) | 0.031* |
| **Right pulvinar → Right amygdala** | 13017±12151.53 | 7829.67±13305.86 | 1.66 (1.64 – 1.69) | <0.001*** |
| **Right pulvinar → Right pSTS** | 1543±1383.55 | 4057.67±2831.43 | 0.38 (0.07 – 1.97) | 0.226 |
| **Global Tractography** | | | | |
| **Tract** | **Actual Mean ± SD** | **Null Mean ± SD** | **IRR (95% CI)** | **p-value** |
| **Left pulvinar → Left amygdala** | 639.67±792.37 | 40.67±67.00 | 15.73 (0.95 – 261.65) | 0.030* |
| **Left pulvinar → Left pSTS** | 1.67±1.53 | 36.33±22.14 | 0.05 (0.01 – 0.16) | <0.001*** |
| **Right pulvinar → Right amygdala** | 1185±1991.31 | 58.33±99.31 | 20.31 (17.51 – 23.73) | <0.001*** |
| **Right pulvinar → Right pSTS** | 7.33±6.35 | 56.33±25.72 | 0.13 (0.04 – 0.41) | 0.0004*** |
